# Single-molecule visualization of sequence-specific RNA binding by a designer PPR protein

**DOI:** 10.1101/2024.04.22.590477

**Authors:** Nicholas Marzano, Brady Johnston, Bishnu P. Paudel, Jason Schmidberger, Slobodan Jergic, Till Böcking, Mark Agostino, Ian Small, Antoine M. van Oijen, Charles S. Bond

## Abstract

Pentatricopeptide repeat (PPR) proteins are a large family of modular RNA-binding proteins that recognize specific ssRNA target sequences. There is significant interest in developing ‘designer’ PPRs for use in diagnostics or as tools to detect and localize target RNA sequences. However, it is unclear how PPRs search for target sequences within complex transcriptomes and current models to predict PPR binding sites struggle to reconcile the effects that RNA mismatches and secondary structure have on PPR binding. To address this, we determined the structure of a designer PPR (dPPR10) bound to its target sequence and used two- and three-colour single-molecule FRET to interrogate the mechanism of ssRNA binding to individual PPR proteins in real time. We demonstrate that longer RNA sequences were significantly slower to bind (or could not bind at all) and that this is, in part, due to their propensity to form stable secondary structures that sequester the target sequence from dPPR10. Importantly, dPPR10 does not associate with non-target flanking sequences, binding specifically to its target sequence within longer ssRNA species. This data provides evidence that PPRs have limited to no capacity to ‘scan’ RNA transcripts for target sequences and instead rely on diffusion for cognate searching. The kinetic constraints imposed by random three-dimensional diffusion may explain the long-standing conundrum of why PPR proteins are abundant in organelles, but almost unknown outside them (i.e. in the cytosol and nucleus). These findings will inform improved prediction of PPR binding sites for the development of designer PPRs.

**Summary:** Pentatricopeptide repeat proteins (PPR) are a large family of modular RNA-binding proteins, whereby each module can be ‘designed’ to bind to a specific ssRNA nucleobase and thus any RNA sequence of interest. As such, there is substantial interest in developing ‘designer’ PPRs for a range of biotechnology applications, including diagnostics or *in vivo* localisation of RNA species; however, the mechanistic details regarding how PPRs search for and bind to target sequences is unclear. As such, we combined structure-based and single-molecule approaches and determined that PPRs bind only to their target sequences (i.e., they do not associate with non-target RNA sequences) and do not ‘scan’ longer RNA oligonucleotides for the target sequence. Instead, target searching appears kinetically-constrained by random three-dimensional diffusion, providing an explanation as to why PPRs are found almost exclusively in organelle compartments that typically have smaller transcriptomes. Collectively, this work identifies several key considerations for future ‘designer’ PPR developments.

## Introduction

It is increasingly apparent that the plethora of RNA forms (e.g., mRNA, long non-coding RNA, spliceosome components) play complex roles in information transfer and regulation within cells ^1^. Indeed, the functional output of even a single RNA transcript can vary depending on several factors, such as RNA stability, secondary structure or splicing variations, many of which are regulated by classes of RNA-binding proteins. One of these classes are the pentatricopeptide repeat (PPR) proteins, which are a large family of modular RNA-binding proteins with many roles in RNA stability, processing, splicing, and translation ^2–7^. While prevalent in all eukaryotes, the PPR family is extensively expanded in terrestrial plants and its members are almost uniquely found in organelles ^8^. PPR proteins bind to specific single-stranded RNA (ssRNA) target sequences; their specificity is ultimately determined by repeating modular motifs of *ca.* 35 amino acids, whereby each consecutive module specifically recognises a discrete RNA nucleobase. A PPR recognition code ^9,10^ relates how each nucleotide interacts with two amino acids at the 5^th^ and final (commonly 35^th^) positions of each motif; amino acid 5 primarily distinguishes purines from pyrimidines and amino acid 35 primarily distinguishes between guanine/uracil or cytosine/adenine bases. The determination of the PPR recognition code has resulted in the emergence of ‘designer’ PPR (dPPR) proteins, which can be rationally modified to target any RNA sequence of interest ^11–14^. As such, there is significant potential for the use of dPPRs in diagnostics (e.g., RNA detection in biological fluids), as a tool to modulate the expression or silencing of target genes ^15–17^, or to detect and localize target RNA sequences *in vivo.* However, to be effective, the affinity of a dPPR to its binding site needs to be sufficiently high, yet simultaneously specific to minimise the occurrence of off-target binding.

While the PPR code is a useful tool to help predict PPR binding sites, there is a two-fold degeneracy that often makes the prediction of both PPR binding sites and the sequence specificity of naturally occurring PPR proteins *in vivo* ambiguous ^18,19^. This degeneracy of the PPR code occurs since multiple amino acid combinations can specify for the same nucleotide base, and conversely, the same amino acid combinations can target a variety of nucleotides. Additionally, not all amino acid combinations bind to nucleotides with equal affinity ^20^ and the importance of nucleotides for recognition and binding may vary according to their position within the binding site ^18,21^. Target prediction becomes even more complex in the context of the cellular environment, where PPR protein target sites often contain ‘mismatches’, or are located within long RNA sequences that contain substantial folded secondary structure. While PPR proteins can prevent the formation of RNA secondary structure ^7^, it remains unclear how transcript length affects the kinetics of PPR binding. Critically, the precise manner by which PPRs search, recognize and bind to their target sequence within much longer transcripts is not known; for example, do PPRs bind to the end of a transcript and ‘scan’ for the target sequence, or do they bind directly to the target (with the ensuing topological issues)? Understanding these fundamental aspects of PPR function may help explain why PPR proteins are predominantly restricted to organelle compartments, which has been a long-standing conundrum in the field. As such, we sought to investigate the precise mechanism(s) by which PPR proteins bind to their target sequences and are challenged by factors present *in vivo* (e.g., mismatches, longer transcripts, etc.) to ultimately improve the prediction of binding sites and the design of synthetic PPR proteins for *in vivo* and diagnostic applications.

To do so, we have designed a synthetic dPPR protein that is a mimic of the model protein *Arabidopsis thaliana* PPR10, such that it is targeted to a 17-nucleotide region of the transcript *atpH* ^7^, and investigated its binding to a range of RNA variants using a combination of X-ray crystallography, microscale thermophoresis, surface plasmon resonance and single-molecule fluorescence approaches. We present the first reported crystal structure of a designer P-class PPR protein (dPPR10) bound to a complex target RNA molecule, in which we observe that dPPR10 becomes conformationally compressed when bound to RNA. Using this information to design and validate a sequence-specific RNA-stimulated FRET-capable protein, we then performed two- and three-colour single-molecule fluorescence resonance energy transfer (smFRET) experiments that allow the binding of ssRNA constructs to a single dPPR10 protein to be monitored in real time. We confirm that mismatches toward the center of the binding site, but not the 3’ end, of the cognate sequence are poorly tolerated and that RNA secondary structure affects the accessibility of binding sites to dPPR10. In addition, we demonstrate that dPPR10 does not associate with non-cognate sequences and instead binds directly to the target, suggesting that it has little or no ability to scan RNA transcripts. Together, this implies there would be kinetic constraints on target recognition in complex transcriptomes that would explain why PPR proteins are primarily localised to organelles and not found more widely distributed throughout the cellular milieu.

## Methods Materials

### Nucleic acids

Unmodified or fluorescently-labelled RNA and DNA oligonucleotides were purchased from Integrated DNA Technologies (IDT) at the 100 nmol scale. All sequences are provided in Figure S1A. Oligonucleotides were resuspended in RNAse-free water to a concentration of 1 mM and stored at -80°C until further use.

### Generation and confirmation of RNA-DNA duplexes

RNA-primer duplexes were generated by incubating RNA (20 µM) in the absence or presence of increasing molar ratios of DNA primer (1:1 – 1:2 RNA:primer) in annealing buffer (25 mM Tris [pH 7.8], 50 mM KCl). Samples were heated at 95°C for 2 min to resolve any secondary structure or nucleotide dimers and then cooled to 4°C by incremental decreases in temperature (1°C every 38 s, Mastercycler Nexus GX2). DNA primer in the absence of RNA was included as a control. To confirm successful formation of RNA-primer duplexes, the samples were diluted 1:20, loaded onto a 4-20% Mini-PROTEAN TGX gel (BioRad) and run in TAE buffer (40 mM Tris, 20 mM acetic acid and 1 mM EDTA) at 100 V for 50 min. The gel was then treated in fixing buffer (10% [v/v] methanol, 7% [v/v] acetic acid and 5% [v/v] glycerol) for 15 min, washed 3x in milli-Q water and then stained with 1x SYBR-Gold in TAE buffer for 15 min. The gel was then imaged using a Typhoon imager as per the manufacturer’s instructions. Samples containing complete RNA-primer duplexes were then stored at -80°C until use.

### Protein expression, purification and chemical labelling

The design of the dPPR10 construct followed the methodology put forward in Gully *et al 2015* ^11^, which involved repeating a consensus 35-amino-acid repeat sequence 17 times, changing residues 5 and 35 of the motif to target it to each base of the target RNA. The final amino acid sequence of purified, cleaved dPPR10 protein is GAMGND-(VVTY[<u>NT</u>]TLIDGLAKAGRLEEALQLFQEMKEKGVKP[<u>DNS</u>])_17_-VVTNNTLKDGASKAG, where the bracketed motif is repeated, and the square-bracketed underlined residues that vary (Figure S2). The sequence of the variant protein dPPR10-C2 is identical to dPPR10 except for the substitution of residues Gln96 and Gln376 to cysteine, and the inclusion of an N-terminal Avi-tag sequence for biotinylation (Figure S2). The complete protein sequences for dPPR10 and dPPR10-C2 are given in Table S1. Genes were codon-optimised, flanked with NcoI and EcoRI restriction endonuclease sites, synthesized and delivered in a pUC57 vector by GenScript. The insert was subsequently subcloned into pETM20 (a gift from Gunter Stier, EMBL Protein Core Facility, Heidelberg, Germany; https://www.embl.de) for protein overexpression and purification. Transformed colonies were plated on LB agar plates supplemented with ampicillin (100 µg/mL) for selection and maintenance of the pETM20 plasmid.

The dPPR10 protein was overexpressed in Rosetta 2™ (DE3) *Escherichia coli*, following the method of Gully *et al* (2015) ^11^ with minor modifications. LB medium (0.5 L) was innoculated with transformed cells and incubated with shaking at 180 rpm at 37°C until the culture reached an optical density of 0.6. Expression was induced with 0.5 mM IPTG, then shaken at 16°C for 18 h.

For biotin-mediated immobilisation of dPPR10-C2, the Avi-tag/BirA coexpression method was used ^22^. Vector pETM20-avi-dPPR10-C2 was co-expressed in BL21(DE3) with pCY216, which encodes the biotin ligase BirA under control of an arabinose promoter (a gift of Steven Polyak, University of Adelaide). When cells, cultured in LB medium supplemented with a final concentration of 0.2 mg/mL biotin, reached an optical density of 0.6, dPPR10-C2 and BirA were both induced upon addition of 0.5 mM IPTG and 0.05% (w/v) L-arabinose and incubated overnight at 16°C at 180 rpm for 18 h.

Cells were harvested by centrifugation at 4,000 x g for 10 min with pellets frozen and stored at -20°C. Pellets were resuspended in 50 mM Tris-HCl (pH 8.0), 500 mM KCl, 10% (v/v) glycerol (lysis buffer) with a mini cOmplete protease inhibitor (EDTA free) tablet (Roche) and 0.5 µL Benzonase 250 U/µL (Sigma Aldrich). Bacteria were lysed under high pressure using an Emulsiflex C5 Homogeniser (Avestin). The insoluble fraction was pelleted by centrifugation at 24,000 g for 30 min at 4°C. The supernatant containing soluble proteins was loaded on to a 5 mL HisTrap Column (GE Healthcare) at 2.5 mL/min and washed at 2.5 mL/min with 10% elution buffer (lysis buffer + 500 mM imidazole). Protein was eluted from the column using a 10-100% gradient of elution buffer over eight column volumes. Protein-containing fractions were pooled and dialysed overnight into lysis buffer in the presence of 4 mg TEV protease and DTT (1 mM). The cleaved protein was washed through a 5 mL HisTrap column to remove any remaining His-tagged protein impurities. SDS-PAGE evaluated protein purity (Mini-PROTEAN TGX Stain-Free Gels, 4-20% 15-well comb, Biorad). Pure protein fractions were typically concentrated to 12-14 mg/mL using centrifugal concentration (Amicon™ Ultra – 30 kDa molecular weight cut-off [MWCO], Merck Millipore) before being snap-frozen in liquid nitrogen and stored at -80°C.

### Protein Labelling with Fluorescent Dyes

dPPR10-C2 was fluorescently labelled with a Cy3 and Alexa Fluor 647 (AF647) FRET-pair as described previously ^23^ with minor modifications. Briefly, dPPR10-C2 (∼ 2 mg/mL) was incubated in the presence of 5 mM Tris(2- carboxyethyl)phosphine (TCEP) and placed on a rotator for 1 h at 4°C. dPPR10 was then incubated in the presence of a 4-fold and 6-fold excess of pre-mixed Cy3 donor and AF647 acceptor fluorophores, respectively, and placed on a rotator overnight at 4°C. Following the coupling reaction, excess dye was removed by gel filtration chromatography using a 7K MWCO Zeba Spin Desalting column (Thermo Fisher Scientific, USA) equilibrated in 50 mM Tris (pH 7.5) supplemented with 20% (v/v) glycerol. The concentration and degree of labelling were calculated by UV absorbance.

### Structure determination and biophysical characterization of dPPR10

#### Analytical Size Exclusion Chromatography

Protein, RNA and protein-RNA samples were made up to total volume of 25 µL in 50 mM Tris-HCL (pH 8.0), 100 mM KCl. Protein was diluted to 1 mg/mL (15 µM), with RNA added at a final molar ratio of 1:1.2. A volume of 12.5 µL of sample was injected onto a Superdex 200 increase 5/150 column (Cytiva), at a flow rate of 0.3 mL/min. Absorbance was measured at 280 and 260 nm using a Biologic Quadtech UV/VIS detector (Biorad).

#### Crystallographic Structure Determination

Concentrated (12.4 mg/mL) dPPR10 was incubated with its target ssRNA sequence (denoted as RNA^cognate^, Figure S1A) at a 1:1.2 molar ratio and applied to several commercially available 96-well crystallisation screens using an Art Robbins Phoenix, SBS-format Intelliplates and drops formed from 100 nL protein and 100 nL reservoir. Diffraction-quality crystals grew in 100 mM magnesium acetate, 50 mM MES (pH 5.6), and 20% 2- methyl-2,4-pentanediol through sitting drop vapor diffusion.

Single crystals with dimensions of *ca.* 100 µm were harvested directly from the 96 well plate in a nylon loop (Hampton Research), flash frozen in liquid nitrogen and screened for diffraction at the MX2 beamline of the Australian Synchrotron ^24,25^ . A complete dataset to 2.0 Å resolution was collected on a Dectris Eiger 16M detector, at 100 K, with 0.1° *ϕ* rotations, 180° wedge collected over 18 s with 60% beam attenuation and a crystal-to-detector distance of 315 mm.

Diffraction data were indexed, merged and integrated using XDS ^26^. Subsequent processing used the CCP4 suite ^27^. Scaling and space group identification was performed using AIMLESS ^28^, and the structure was solved by molecular replacement using PHENIX ^29^ with a search model modified from PDB 5I9F using a single protein chain containing eight PPR motifs ^12^. Model building was completed using iterative cycles of graphical building in COOT ^30^ and refinement in REFMAC ^31^ cycles. The refined structure was deposited at the Protein Data Bank ^32^ with code 6EEN.

#### Surface Plasmon Resonance (SPR) Measurements

Binding studies were performed in SPR buffer (30 mM Tris [pH 7.6], 40 mM KCl, 5 mM MgCl_2_, 0.5 mM DTT, 0.25 mM EDTA and 0.01 % (v/v) P20 surfactant). Temperature was kept constant at 20°C. Biotinylated dPPR10- C2 was immobilized onto the surface of streptavidin-coated (SA) SPR chip by injecting 50 nM dPPR10-C2 at a flow rate of 5 µL/min for 750 s. This yielded *ca.* 1650 response units (RUs) of dPPR10 on the surface. Successive injections of 1 M MgCl_2_ and subsequently 2/3/4 M MgCl_2_ resulted in the loss of some non-specifically bound dPPR10s from the surface, leaving ∼ 1300-1500 RUs of immobilized dPPR10. Given the molecular weight of protein (∼ 70 kDa) and RNA species (∼ 5 kDa), in the case of a 1:1 binding, the response is expected to increase by ∼ 100 RUs upon saturation with RNA (R_max_). The association of RNA from solution to immobilized dPPR10- C2 was monitored following injection of RNA (1 µM) onto the chip at a flow rate of 40 µL/min for 300 s, while its removal is followed by monitoring dissociation over 1000 s.

#### Solution FRET assay

Fluorescently-labelled dPPR10-C2 was thawed and diluted to a final concentration of 5 nM in 100 mM Tris-HCl (pH 8.0), 100 mM KCl. RNA^cognate^ or a ssRNA sequence that was not predicted to bind to dPPR10-C2 based on the PPR code (denoted as RNA^anti-cognate^; Figure S1A) was diluted in a twelve-point concentration series from 1000 nM to 0.01 nM. A variable mode scanner (Amersham Typhoon™, GE Healthcare) was used to measure fluorescence in 96-well microplate, µClear®, Black, non-binding (Greiner Bio-One) with images acquired at a 100 µm per pixel resolution. Fluorescence was recorded on two photon-multiplier tubes with voltages set at 600V. FRET was calculated from the donor (excitation at 532 nm, emission between 560–580 nm) and transfer (excitation at 532 nm, emission between 655–685 nm) channels, following Equation 1.

#### Microscale Thermophoresis (MST)

MST experiments used a Monolith NT 115 Series (NanoTemper Technologies) using 20% LED and 40% IR-laser power for all experiments. Laser on and off times were 30 s and 5 s, respectively. The dilution series (typically 1:1) was prepared by serially diluting a sample of dPPR10 (10 µg/mL [i.e., 150 nM]) in 100 mM Tris-HCl (pH 8.0) and 100 mM KCl, which was then mixed with 5ʹ 6-FAM labelled RNA oligonucleotides (IDT; RNA^cognate^ or RNA^anti-cognate^) to a final RNA concentration of 50 nM. Protein concentration limits were adjusted based on initial experiments to create a complete binding curve. Data was processed and binding curves calculated using MO Affnity Analysis 2.2.7.6056 (NanoTemper).

### Total internal reflection fluorescence (TIRF) microscopy

#### Microscope setup

Samples were imaged using a custom-built TIRF microcopy system constructed using an inverted optical microscope (Nikon Eclipse TI) that was coupled to an electron-multiplied charge-coupled device (EMCDD) camera (Andor iXon Life 897, Oxford Instruments, UK). The camera was integrated to operate in an objective-type TIRF setup with diode-pumped solid-state lasers (200 mW Sapphire; Coherent, USA or Stradus 637-140, Vortran Laser Technology, USA) emitting circularly polarized laser radiation of either 488, 532 or 647-nm continuous wavelength. The laser excitation light was reflected by a dichroic mirror (ZT405/488/532/640; Semrock, USA) and directed through an oil-immersion objective lens (CFI Apochromate TIRF Series 60x objective lens, numerical aperture = 1.49) and onto the sample. Total internal reflection was achieved by directing the incident ray onto the sample at the critical angle (θ_c_) of ∼ 67° for a glass/water interface. The evanescent light field generated selectively excites the surface-immobilized fluorophores, with the fluorescence emission passing through the same objective lens and filtered by the same dichroic mirror. For two-colour experiments, the emission was then split using a T635lpxr-UF2 dichroic (Chroma, USA), passed through ET690/50m and ET595/50m (Chroma, USA) cleanup filters and the final fluorescent image projected onto the EMCDD camera. For three-colour experiments, the emission was split using T635lpxr-UF2 and T550lpxr-UF2 dichroics, passed through ET690/50m, ET595/50m and ET525/50 cleanup filters and the final fluorescent image projected onto the EMCDD camera. The camera was running in frame transfer mode at 5Hz, with an electron multiplication gain of 700, operating at -70°C with a pixel distance of 160 nm (in sample space).

#### Coverslip preparation and flow cell assembly

Coverslips were functionalized with neutravidin as previously described ^33^, with minor modifications. Briefly, 24 x 24 mm glass coverslips were cleaned by alternatively sonicating in 100% ethanol and 5 M KOH for a total of 2 h before aminosilanization in 2% (v/v) 3-aminopropyl trimethoxysilane (Alfa Aesar, USA) for 15 min. NHS-ester methoxy-polyethylene glycol, molecular weight 5 kDa (mPEG) and biotinylated-mPEG (bPEG; LaysanBio, USA), at a 20:1 (w/w) ratio, was dissolved in 50 mM 4-morpholinepropanesulfonic acid (MOPS, pH 7.4) buffer and sandwiched between two activated coverslips for a minimum of 4 h for initial passivation in a custom-made humidity chamber. PEGylated coverslips were then rinsed with milli-Q and PEGylated again as described above for overnight (∼ 20 h) passivation. PEGylated coverslips were rinsed with milli-Q water, dried under nitrogen gas, and stored at -20°C until required. Prior to use, neutravidin (0.2 mg/mL; BioLabs, USA) in milli-Q was incubated on the passivated coverslip for 10 min to bind to the bPEG. Neutravidin functionalized coverslips were then rinsed with milli-Q, dried under nitrogen gas and bonded to a polydimethylsiloxane (PDMS) flow cell for use in single-molecule experiments. Finally, to reduce the non-specific binding of proteins to the coverslip surface, each channel in the microfluidic setup was incubated in the presence of 2% (v/v) Tween-20 for 30 min as previously described ^34^ and then washed with imaging buffer.

### smFRET experiments and analysis

#### Surface immobilization of labelled proteins and acquisition of smFRET data

For smFRET experiments, fluorescently-labelled dPPR10-C2 was specifically immobilized to a neutravidin-functionalized and Tween-20-coated coverslip. To do so, labelled protein (∼ 50 pM final concentration) was diluted in imaging buffer (50 mM Tris [pH 8.0], 250 mM KCl and 6 mM 6-hydroxy-2,5,7,8-tetramethylchroman- 2-carboxylic acid [Trolox]) and incubated in the flow cell for 5 min. These conditions would typically give rise to ∼ 75–100 FRET-competent molecules per 100 x 100 µm. Unbound proteins were then removed from the flow cell by flowing through imaging buffer.

For two-color experiments, data was acquired using the TIRF microscope setup previously described following sample illumination using a 532-nm solid state laser with excitation intensity of 2.6 W/cm^2^ and the fluorescence of donor and acceptor fluorophores was measured every 200 ms at multiple fields of view. For three-color experiments, the sample was alternatively illuminated using a 532-nm and 488-nm solid state for 200 ms per frame and the fluorescence of all dyes measured. An oxygen scavenging system (OSS) consisting of 5 mM protocatechuic acid and 50 nM protocatechuate-3,4-dioxygenase was included in all buffers prior to and during image acquisition to minimize photobleaching and fluorophore blinking.

To monitor dPPR10-C2 conformational changes upon incubation with ssRNA constructs, a combination of live- flow and steady state smFRET experiments were performed. For live-flow conditions, imaging was initiated and ssRNA (1 µM) was injected into the microfludic flow cell using a syringe pump (300 µL, NE-1000, ProSense B.V) after 10 s; imaging continued for 6 min to observe subsequent changes in FRET efficiency. Multiple fields of view were subequently imaged for 6 min each in the presence of the incubated ssRNA and the data collated (i.e., steady-state conditions, minimum of 30 min). Three-color experiments were performed as described above, with the exception that the concentration of fluorescently labelled ssRNA (20 nM) was reduced to prevent fluorescence background.

#### Molecule selection and FRET calculations

Single-molecule time trajectories were analyzed in MATLAB using the MASH-FRET user interface (version 1.2.2, accessible at https://rna-fretools.github.io/MASH-FRET/) ^35^. The approximate FRET value is measured as the ratio between the acceptor fluorescence intensity (*I_Acceptor_*) and the sum of both donor (*I_Donor_*) and acceptor fluorescence intensities after correcting for crosstalk between donor and acceptor channels. The formula for calculating the FRET efficiency is given by Equation 1, whereby the corrected acceptor intensity (denoted as *CI_Acceptor_*) is equal to *I_Acceptor_ –* (g * *I_Donor_*) and g is the crosstalk correction constant. g is calculated as the ratio of fluorescence measured in the acceptor and donor detection channels following direct excitation of a protein labelled with a single donor fluorophore.

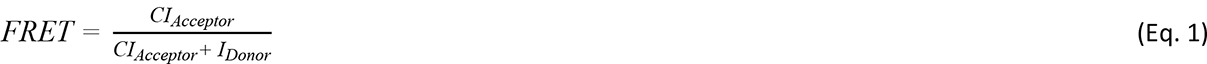

Briefly, donor and acceptor fluorescence channels were aligned following a local weighted mean transformation of images of TetraSpeck fluorescence beads and donor and acceptor fluorescence spots co- localized to identify FRET pairs. Molecules that displayed clear donor and/or acceptor photobleaching events or demonstrated anti-correlated changes in donor and acceptor fluorescence intensity were selected for subsequent analysis. The number of photobleaching events observed was used to determine the number of fluorophores present; only molecules in which a single donor and acceptor photobleaching event was observed were used for further analysis.

#### Trace processing and Hidden Markov Model (HMM) fitting

Selected molecules were denoised in MASH-FRET using the NL filter, which has been described previously ^36^, to accurately identify and quantify transitions between different FRET states during downstream processing. Parameter values were as follows; exponent factor for predictor weight, 5: running average window size, 1: factor for predictor average window sizes, 2. Data were truncated to only include FRET values acquired before donor or acceptor photobleaching. FRET efficiency data were exported to the state finding algorithm vbFRET (version vbFRET_nov12, https://sourceforge.net/projects/vbFRET/) and trajectories fit to a Hidden Markov Model (HMM) to identify discrete FRET states, their residence times, and the transition distributions between them. Default vbFRET settings were employed to fit data to the HMM, with the exception that the *mu* and *beta* hyperparameters were changed to 1.5 and 0.5, respectively, to prevent over-fitting. For three-colour experiments, the FRET traces were fit with an HMM using custom-written scripts in Python.

#### Kinetic analysis of HMM fits

The HMM fits of individual FRET trajectories were further analyzed to extract key kinetic information arising from changes in PPR conformation. To investigate transitions of interest, each transition (as determined from the HMM analysis) was sorted into different directional classes denoted generally as T_Before-After_, whereby ‘before’ refers to *F*_Before_ and ‘after’ refers to *F*_After_. For simplicity, FRET data were binned according to whether *F*_Before_ or *F*_After_ was greater than 0.5 (high) or less than 0.5 (low), unless otherwise indicated, and thus two different transition classes are possible: T_high-low_ and T_low-high_. The residence time (defined as the time that a molecule resides at *F*_Before_ prior to transition to *F*_After_) for each transition within a transition class was calculated and presented cumulative histograms. Since the FRET data could be well described as a two-state system, the cumulative residence times (determined as the period of time that the HMM fit resides at low-FRET states prior to a T_low-high_ transition, and vice-versa for T_high-low_ transitions) are presented. Since it is not possible to determine for how long a particular FRET state would have existed if not truncated due to photobleaching, the last measured FRET state was deleted and thus excluded from residence time calculations. Since data were smoothed during denoising, residence times shorter than that given by Equation 2 were not considered for further analysis. *F* is the imaging frame rate in milliseconds and *N_FA_* is the number of frames that were averaged during trace denoising.

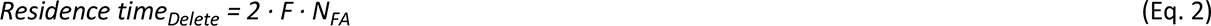

The rate of ssRNA binding-and-release events was determined by calculating the number of transitions to FRET states below (i.e., release event) or above (i.e., binding event) the FRET threshold set for ssRNA binding (0.5 FRET) for each molecule. From here, the minimum number of binding or release events was divided by the imaging lifetime of that molecule (i.e., time until photobleaching) to determine the binding-and-release rate. Finally, to determine whether there are changes in FRET immediately prior to ssRNA binding, all T_low-high_ transitions were identified (which are indicative of ssRNA binding), filtered such that only transitions with a residence time > 10 s were selected, and the FRET efficiency 10 s prior to and after each T_low-high_ was plotted.

The FRET efficiency of all datapoints for each molecule was collated and are presented as FRET efficiency histograms.

### Statistical analysis

To determine how different ssRNA constructs affect the lifetime of certain FRET states, the residence times for each transition class were statistically analyzed using a one-way analysis of variance (ANOVA) with a Tukey’s multiple comparisons post-hoc test, with p ≤ 0.05 determined to be statistically significant. All data analysis and presentation were performed using custom-written scripts on Python software or using GraphPad Prism 9 (GraphPad Software Inc; San Diego, USA). All original code for the analysis and visualization of smFRET data has been deposited at Zenodo at the following DOI: 10.5281/zenodo.10989968.

## Results

### Designer PPR proteins undergo a predictable, uniform conformational change upon binding RNA

The complex of dPPR10 bound to its cognate target (RNA^cognate^; including the 17 nt fragment of *atpH* RNA) was readily crystallised in spacegroup *P1* (unit cell: 43.35 Å, 51.65 Å, 51.97 Å, 118.12°, 97.21°, 96.00°), yielding a structure to 2.0 Å resolution (*R* = 0.183, *R_free_* = 0.243, Figure 1A, Table S2). While the protein-RNA complex contains 17 PPR repeats, the crystal structure reflects a continuous protein-RNA superhelical complex (Figure S3A) with 9 repeats per superhelical turn (Figure 1A). Helical disorder observed in this structure is of the same form reported previously for tetratricopeptide repeat and PPR proteins ^11,37^, resulting in a structure with excellent electron density and geometry for an extensive protein-RNA molecule that spans the crystal (Figure 1B, Figure S3A). Microheterogeneity at the nucleobases and the base-recognising amino acid sidechains in positions 5 and 35 of each repeat results in averaged electron density for these regions (Figure S3B).

**Figure 1:**
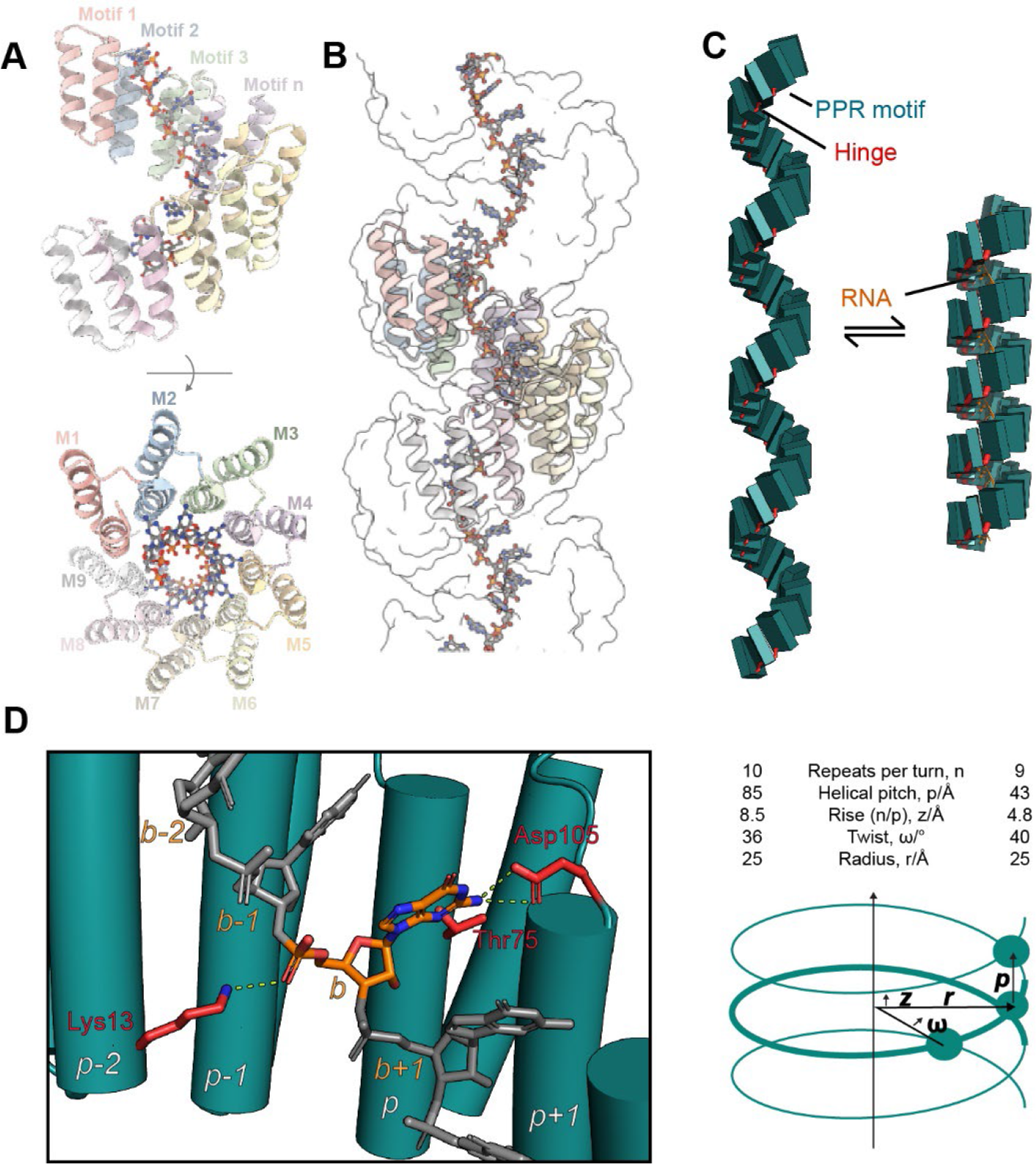
Structural parameters of conformational change of dPPR10 on binding its target RNA. (A) Cartoon representation of orthogonal views of the dPPR10 crystal unit cell contents. The individual PPR motifs are denoted in text. **(B)** Neighbouring unit cells (shown as surface representation) result in an extended idealised superhelical protein:RNA complex structure. **(C)** Comparison of dPPR (PDB 4OZS) and dPPR10:RNA (PDB 6EEN) structural parameters reveals a dramatic, uniform conformational change on RNA binding (boxes represent individual PPR motifs). **(D)** Each nucleotide interacts with amino acids (e.g., Lys13 from *p-2*, Thr75 and Asp105 from motif *p* for nucleobase *b*) from three adjacent PPR motifs that act to stabilize the bound state. Note that Asp105 is the last residue of motif *p* and directly precedes the start of motif *p+1*. Such interactions likely do not contribute to PPR-RNA specificity. Figure prepared using PYMOL (The PyMOL Molecular Graphics System, Version 2.0 Schrödinger, LLC.), PDB-MODE ^40^ and Blender (Community, B. O. (2018). Blender - a 3D modelling and rendering package. Stichting Blender Foundation, Amsterdam. Retrieved from http://www.blender.org) with Molecular Nodes plugin (Johnston, 2022 “MolecularNodes”. Retrieved from https://bradyajohnston.github.io/MolecularNodes/)

Both the dPPR10:*atpH* structure and the structure of dPPR apo-protein ^11^ (PDB entry 4OZS), which effectively differs from dPPR10 only in the base-interacting amino acids 5 and 35, reveal an idealised model of the PPR α-solenoid superhelix that allows objective measurement and comparison of superhelical parameters (Figure 1C). The dPPR10:*atpH* superhelix is substantially contracted in comparison with the apo protein (Figure 1C), as expected based on previous reports ^11,12,38^. The superhelical pitch, which can be measured directly from the lattice parameters, is almost halved to 43 Å (*cf.* 85 Å for apo-protein), and is overwound, with nine repeats per helical turn (*cf.* ten). Using the parameters defined previously ^39^, this corresponds to a rise of 4.8 Å (*cf.* 8.5 Å), a twist of 40° (*cf.* 36 Å) and a consistent superhelical radius of ∼ 25 Å.

To gain a better understanding of the mechanism of conformational change on RNA binding, we performed a detailed geometric analysis. Initial inspection showed that the local structural changes are surprisingly subtle relative to the substantial compaction of the superhelix. The program DYNDOM ^41^ reveals that individual PPR repeats can be approximated as rigid bodies (RMSD 0.4-0.6 Å), which rotate with respect to each other around a contiguous hinge comprised of residues 33-35 at the junction of adjacent motifs (Figure 1C; Figure S3C).

Rotation of 12° ± 4° and translation of up to 1.2 Å about this hinge allows interconversion between the extended and compact structures. Close investigation of backbone geometry (Figure S3D) shows that apart from the flexible hinges, the individual motifs are relatively rigid. However, notable torsion occurs at residues 13, 16-17, 27 and 32-35. Of these residues, 32-35 are involved in the hinge, while Met27 is a highly conserved residue which interacts directly with the hinge residues. Residues 16-17 comprise the two-residue linker between helices within a PPR repeat and may thus be expected to absorb some torsional strain. Experimentally, the contraction of dPPR10 upon binding RNA^cognate^ is readily observed in size-exclusion chromatography, where the complex runs slower than the apo-protein despite having a larger mass, due to its compact hydrodynamic profile (Figure S4A).

Notably, Lys13 is a highly conserved RNA-interacting residue, which may have importance linking RNA binding to the change in orientation of PPR repeats. Any given RNA nucleotide binds a span of three consecutive PPR repeats (Figure 1D), such that (i) nucleobase *b* is recognised by sidechains of residues 5 and 35 from PPR motif *p* and, (ii) the 5’-phosphate group of nucleobase *b* is bound by a salt-bridge from Lys13 from PPR motif *p*-2. As the mean distance between adjacent Lys13-Nζ atoms reduces from 9.2 ± 2.7 Å to 6.7 ± 0.2 Å on binding to RNA, it is likely that coulombic repulsion between these lysine residues contributes to the more extended apoprotein structure (Figure S3E). Taken together, this implies that the sequential binding of nucleotides proceeds via conformational changes to overlapping three-motif segments of the PPR protein.

### Binding of RNA by designer PPR proteins can be detected by conformational change-induced increases in intramolecular FRET

Using our idealised RNA-bound and protein-only structures, we comprehensively evaluated the geometric relationship of pairs of residues to select which residues if suitably labelled would span the Förster critical transfer distance for commonly used FRET pairs. We identified Gln22 of PPR motifs 8 repeats (280 residues) apart as being potentially suitable for this purpose; these residues are polar, located on the external face of the superhelix, and have an inter-residue distance of 71 Å in the absence of RNA, and 40 Å in its presence, which lie on either side of the Förster critical transfer distance (54 Å) for a Cy3/Alexa Fluor 647 (AF647) FRET pair (Figure 2A). Substitution of these residues in repeats 3 and 11 to cysteine (Figure S2) and optimisation of a dual-labelling reaction with maleimide-functionalised dyes resulted in a sample in which approximately 50% of the protein is labelled with both dyes. In a plate based fluoresence assay, dPPR10 showed concentration-dependent changes in acceptor fluorescence in the presence of the RNA^cognate^ oligonucleotide commensurate with a dissociation constant of 1.2 nM (Figure S4B-C), confirming the suitability of replacing Gln22 residues with dye-attaching Cys residues. This binding affinity compares favourably to the 4 nM dissociation constant determined by microscale thermophoresis (MST) for unsubstituted, unlabelled dPPR10 protein binding to

**Figure 2:**
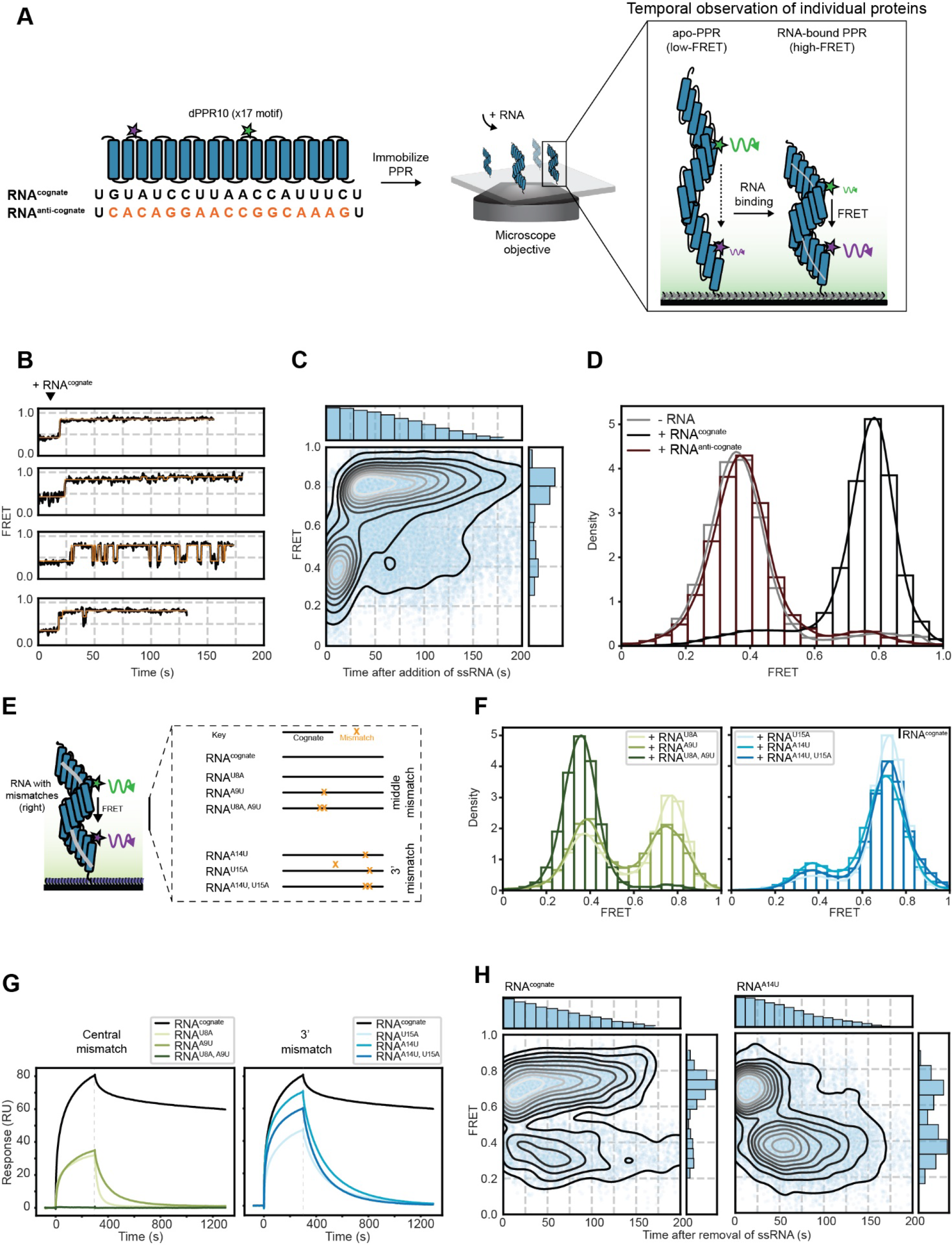
smFRET can be used to monitor dPPR10 conformational changes upon association or dissociation of target and mismatched ssRNA oligonucleotides. (A) Schematic of the dPPR10 protein and imaging setup used for smFRET experiments. The architecture of dPPR10 shows the alignment of each motif with its corresponding nucleobase in the cognate sequence (RNA^cognate^) and approximate location of attached fluorophores used for smFRET (*left*). Note that the first and last nucleotide of RNA^cognate^ does not pair with a PPR motif. The Cy3/AF647-labelled dPPR10 protein is immobilized to a coverslip surface and the fluorescence of both dyes are measured and used to determine the FRET efficiency of protein in the absence or presence of ssRNA (*right*). **(B)** Representative FRET trajectories of dPPR10 upon injection of RNA^cognate^ (1 µM) after 10 s (indicated above the traces). **(C)** A heatmap of the FRET intensity over time following the injection of RNA^cognate^ as described in D. Histograms show the decrease in data density due to photobleaching over time (*top*) and the collated FRET efficiency (*right*). Data is collated from 168 individual molecules. **(D)** FRET histogram of dPPR10 incubated in the absence or presence of the indicated ssRNA oligonucleotides (1 µM). Data is collated from at least 114 individual molecules. **(E)** Schematic showing the position of different mismatches introduced into RNA^cognate^ for subsequent smFRET experiments. The key denotes the sequences that are identical (*black*) or different (*orange*) to RNA^cognate^. **(F)** FRET histogram of dPPR10 incubated in the absence or presence of the indicated ssRNA oligonucleotides (1 µM). ssRNA sequences containing mismatches in the dPPR10 region probed by FRET (*blue, left*) and those at the 3’ end (*green, right*) is indicated. Data is collated from at least 146 individual molecules. **(G)** Surface plasmon resonance (SPR) traces of ssRNA constructs (1 µM) binding to immobilized dPPR10. Association phase of ssRNA to dPPR10 is initiated at t = 0 and proceeds for 300 s. Dissociation curves are generated following removal of ssRNA from solution. Data for RNA^U8A^, RNA^A9U^ and RNA^U8A, A9U^ (*left*) or RNA^U15A^, RNA^A14U^ and RNA^A14U, U15A^ (*right*) are shown. **(H)** Heatmaps of the FRET intensity over time following the removal of ssRNA (at 10 s) from dPPR10 molecules previously incubated with and bound to RNA^cognate^ (*left*) or RNA^A14U^ (*right*). Data is collated from at least 43 individual molecules.

FITC-labelled RNA^cognate^ (Figure S4D). Both FRET and MST ensemble assays showed that dPPR10 does not bind to the non-cognate RNA^anti-cognate^.

### The binding of cognate or mismatched ssRNA to individual dPPR10 proteins can be monitored temporally using single-molecule FRET

While methods that analyze large populations of molecules in bulk as time- or ensemble-averages (such as crystallography, spectroscopy, thermophoresis, SPR) can be used to reveal important aspects of structure, thermodynamics and kinetics, single-molecule methods have the power to interrogate mechanistic details in ways that other methods simply cannot. We therefore sought to take advantage of the significant conformational rearrangements of dPPR10 upon binding to ssRNA (Figure 1C), and its demonstrated capability as a FRET-based sensor (Figure S4), by developing a single-molecule FRET (smFRET) assay that could be used to monitor dynamic changes in protein conformation in the presence of various ssRNA oligonucleotides. To do so, we generated an enzymatically biotinylated dPPR10 protein (i.e., dPPR10-C2, referred to as dPPR10 henceforth for simplicity) labelled with Cy3 and AF647 that could be specifically immobilised to a coverslip surface via streptavidin-biotin interactions and imaged in the absence or presence of a selection of ssRNA variants (Figure 2A). Using microfluidics, we initiated the binding of ssRNA to dPPR10 by injecting its target ssRNA oligonucleotide (RNA^cognate^) after 10 s of imaging (Figure 2B-C). Initially, in the absence of ssRNA, dPPR10 adopts a stable, low-FRET (∼ 0.4) conformation that is consistent with the expanded α-solenoid structure reported previously in the absence of ssRNA ^11^. Upon the addition of RNA^cognate^, there was a pronounced one-step increase in FRET efficiency from the unbound (∼ 0.4) to the ssRNA-bound (∼ 0.8) conformation of dPPR10. Initial binding of ssRNA was predominantly (80%) observed to be stable (i.e., no dynamic changes in FRET post initial binding) or, in rarer cases (20%), unstable (i.e., multiple FRET transitions are observed post initial binding) (Figure 2B). Importantly, dPPR10 adopted the high-FRET state (∼ 0.8) only when incubated in the presence of RNA^cognate^ and not when incubated in the presence of non-target RNA (i.e., RNA^anti-cognate^) (Figure 2D), demonstrating the specificity of dPPR10 for its target.

We further exploited the smFRET assay to investigate how mismatches at different regions of the ssRNA cognate sequence affect the affinity and specificity of dPPR10 binding. To do this, we introduced single and double nucleotide substitutions into RNA^cognate^ within the region of dPPR10 that is monitored by smFRET (denoted as RNA^U8A^, RNA^A9U^ and RNA^U8A, A9U^, where the number corresponds to the n^th^ nucleotide from the 5’ end) (Figure 2E). When dPPR10 was incubated in the presence of the single-nucleotide mismatched ssRNA, FRET trajectories dynamically transitioned between bound and unbound PPR conformations (Figure S5A) and the FRET distributions were bimodal (Figure 2F). The double mismatched ssRNA (i.e., RNA^U8A, A9U^) was unable to induce any increase in FRET, with the individual traces and histograms appearing like that of dPPR10 in the absence of ssRNA and is consistent with surface plasmon resonance (SPR) data (Figure 2G).

ssRNA constructs were also designed that contained mismatches outside the region of dPPR10 probed by smFRET (denoted as RNA^U15A^, RNA^A14U^ and RNA^A14U, U15A^) (Figure 2E). All three ssRNA constructs were able to induce the high-FRET state characteristic of RNA-bound dPPR10, with the caveat that the absolute FRET efficiency observed (∼ 0.7) was slightly lower compared to dPPR10 in the presence of the ssRNA^cognate^ (Figure 2F). The observed changes within the FRET distributions were consistent with SPR and ensemble-based FRET titrations. RNA species that contained mismatches at the 3’ end were more tolerated by dPPR10 compared to those that contain mismatches centrally within the binding site (Figure 2G, Figure S5B), whereby the amplitude of the SPR signal during the association phase scaled with the height of the high FRET peak in the corresponding single-molecule distributions (Figure 2D and F). Dissociation of RNA^cognate^ measured by SPR was biphasic, revealing a minor fraction of unstable (fast dissociation) and a major fraction of stable complexes (slow dissociation), consistent with the subsets of stable (80%) and dynamic (20%) single-molecule FRET traces observed with RNA^cognate^. Notably, all ssRNA molecules that contained a mismatch formed unstable complexes that dissociated rapidly from dPPR10 when removed from solution, consistent with the frequent switching from the high to the low FRET state in the corresponding single-molecule traces. Since association and dissociation of RNA can occur concurrently during the association phase in SPR, the faster *k*_off_s for mismatched ssRNA appear to mostly explain the lower amplitude observed by SPR during association (note that *k*_on_s were not much perturbed, particularly for RNA^A14U^; Figure S1C). These differences in dissociation kinetics were recapitulated using a combination of microfluidics and smFRET, whereby a decrease in FRET efficiency (indicative of ssRNA dissociation) upon removal of ssRNA from solution was only observed when dPPR10 had been pre-incubated and bound to a mismatched ssRNA (e.g., RNA^A14U^), but not RNA^cognate^ (Figure 2H). Taken together these results indicate that while nucleotides A14 and U15 do not have a critical role in initial binding of RNA to the expanded apo-dPPR10 structure, their relative contribution to binding becomes more prominent as the interaction(s) remodel during solenoid compaction.

To complement the SPR data, we performed kinetic analyses of the individual FRET trajectories by fitting the data to a Hidden Markov Model (HMM). The HMM allows discrete FRET states to be identified within the noise and the duration of each state (denoted as the residence time) to be quantified. For simplicity, FRET states were classified as either low-FRET (< 0.5, corresponding to unbound PPR) or high-FRET (> 0.5, corresponding to ssRNA-bound PPR) and as such, transitions between FRET states can be filtered into two transition classes (high to low, denoted as T_high-low_, and low to high, denoted as T_low-high_). Two of the three ssRNA constructs with middle mismatches (RNA^A9U^ and RNA^U8A, A9U^) had longer T_low-high_ residence times compared to the ssRNA constructs with mismatches in the region of dPPR10 not probed by FRET (i.e., RNA^U15A^, RNA^A14U^ and RNA^A14U, U15A^, since the PPR-motifs that ‘pair’ with these mismatches are more C-terminally located than the two cysteines labeled for FRET) (Figure S1B and S5C). It should be noted that while residence times can be extracted from any observed binding or release event, for some RNA species (e.g., RNA^U8A, A9U^) the occurrence of such transitions are rare (< 20). Regardless, this indicates that mismatches within the FRET-probing region of dPPR10 delay initial binding and/or conformational compaction of dPPR10 and that the 3’ end of ssRNAs are less important for initial recognition and binding to dPPR10 compared to those in the center of the binding sequence. Collectively, this data demonstrates that mismatches detrimentally affect the association of RNA to dPPR10 in a sequence position-dependent manner, and that mismatches drastically decrease the stability of the PPR-RNA complex once RNA is removed from solution (Figure 2G-H).

### Truncated RNA associates faster with dPPR10 while a minimum of 10 PPR-nucleobase contacts are required for conformational compaction

Next, we sought to identify a minimum ssRNA sequence that would induce a conformational change in dPPR10. To do this, we utilized an ssRNA sequence that contained only 8 nucleotides at the 3’ end of RNA^cognate^ (denoted RNA^12–19^) and a series of truncated ssRNA sequences containing the first 10-16 nucleotides at the 5’ end of RNA^cognate^ (denoted RNA^1-10^ to RNA^1–16^) (Figure 3A). When dPPR10 was incubated in the presence of RNA^12–19^, the FRET histogram distributions were like that observed in the absence of ssRNA (Figure 3B), suggesting that dPPR10 is unable to stably interact with this truncated ssRNA target. Similarly, incubation of dPPR10 with the shortest 3’ truncation construct, RNA^1–10^, resulted in FRET distributions similar to dPPR10 in the absence of ssRNA. The addition of a single nucleotide (i.e., RNA^1–11^), however, resulted in individual dPPR10 molecules dynamically transitioning between low-FRET and high-FRET states (Figure S6A) and the appearance of a bimodal FRET distribution (Figure 3B). This data was consistent with SPR results, whereby dPPR10 was not able to bind to RNA^12–19^ but could associate rapidly with RNA^1–11^ (albeit with a lower amplitude than RNA^cognate^); notably, RNA^1–11^ was observed to dissociate rapidly from dPPR10 upon removal of RNA from solution (Figure S6B). As such, RNA^1–11^ represents a minimum ssRNA sequence required to induce conformational compaction of dPPR10, with the subsequent addition of nucleotides to this construct (i.e., RNA^1–12^ to RNA^1–16^) also able to induce stable transitions to the high-FRET state consistent with the ssRNA-bound dPPR10 state. It remains to be tested whether other 11 nucleotide long sections RNA^cognate^ (in the middle or at the 3’ end) can similarly induce compaction, or whether the 5’ region is strictly required. Interestingly, combined incubation of RNA^1–11^ with RNA^12–19^, which together encompasses the entire RNA^cognate^ sequence, resulted in a similar proportion of high-FRET states compared to RNA^1–11^ alone and less than that observed for RNA^cognate^ (Figure S6C). Since association of RNA^12–19^ to dPPR10 is exceedingly slow, the probability that both ssRNA species are bound to dPPR10 is low and as such the simultaneous presence of both sequences (i.e., as is the case for RNA^cognate^) is required for efficient binding. Notably, of the truncation constructs that could induce an increase in FRET, the shortest ssRNA sequence (i.e., RNA^1–11^) induced the highest frequency of switching between FRET states of dPPR10 and this switching frequency was observed to decrease with increasing length of ssRNA (Figure 3C). This is due to most truncated ssRNAs having significantly shorter T_low-high_ and T_high-low_ residence times (i.e., faster binding/dissociation rates) compared to the least truncated ssRNA (Figure S6D), which indicates that shorter ssRNA sequences may have improved accessibility to dPPR10 (but lower stability) compared to longer sequences.

**Figure 3:**
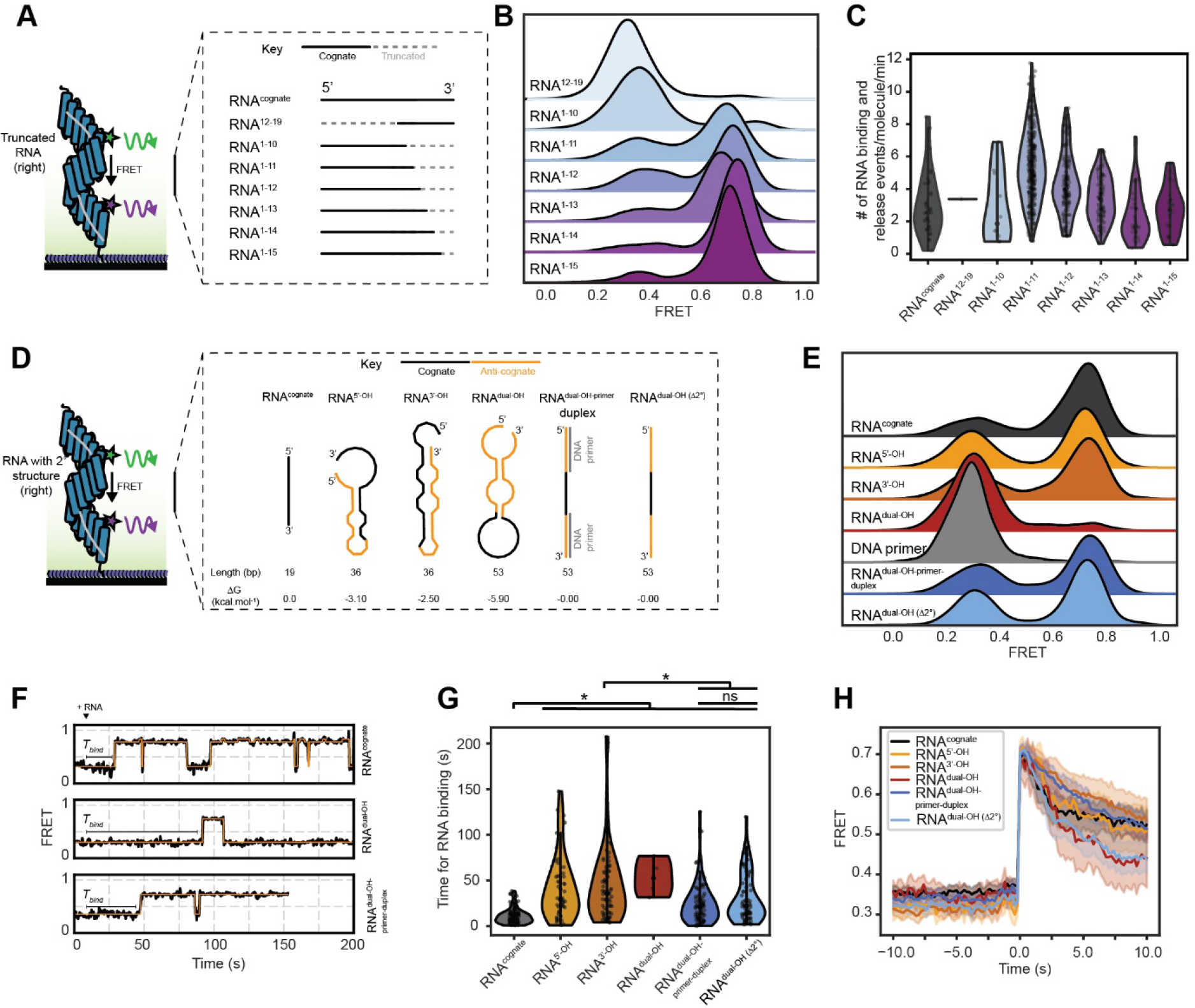
A minimum of 10 PPR-nucleobase contacts are required to induce conformational compaction of dPPR10 while RNA secondary structure regulates access to cognate sequence. (A) Schematic showing the various truncations to RNA^cognate^ that were tested using smFRET. **(B)** Ridgeline plot of the FRET distributions of dPPR10 incubated in the absence or presence of the indicated truncated ssRNA oligonucleotides (1 µM). Data is collated from at least 77 individual molecules. **(C)** Violin plots showing the distribution of ssRNA sampling events by individual dPPR10 molecules in the presence of the different truncated ssRNA oligonucleotides. Note that the rates determined for RNA^cognate^ represents the unstable fraction of complexes that represesent 20% of all traces. **(D)** Schematic of the various extended ssRNA oligonucleotides tested using smFRET and their predicted secondary structure propensities and structures. The position of RNA^cognate^ (*black lines*) and the presence of non-target RNA sequences (*orange lines*) are indicated for each ssRNA. Schematics not shown to scale. **(E)** Ridegeline plot of the FRET distributions of dPPR10 incubated in the absence or presence of the indicated ssRNA oligonucleotides (1 µM). Data is collated from at least 174 individual molecules. **(F)** Representative FRET trajectories of dPPR10 upon the injection of the indicated ssRNA (1 µM) after 10 s (indicated above the traces). The time taken until RNA binding (*T_bind_*) is shown for each trace and used to plot data in panel G. **(G)** Violin plot of the time taken until ssRNA binding. The ssRNA was injected into the flow cell after 10 s and the time until the first transition was determined. A one-way analysis of variance (ANOVA) statistical analysis with Tukey’s multiple-comparisons post-hoc test was performed to determine statistically significant differences in *T_bind_* between treatment groups. *P ≤ 0.05. ns or the absence of markers indicates no significant difference (P > 0.05). **(H)** The average FRET efficiency immediately prior to and after a T_low-high_ transition for dPPR10 in the presence of the indicated ssRNA oligos. T_low-high_ transitions were filtered for those that had residence times longer than 10 s.

### Stable RNA secondary structures sequesters the target sequence and prevent dPPR10 binding

We next sought to investigate how the fusion of flanking non-target sequences to RNA^cognate^ affects the kinetics of dPPR10 binding. To do this, we implemented ssRNA constructs with extensions on either the 5’ or 3’ end (∼ 17 nucleotides, denoted RNA^5’-OH^ or RNA^3’-OH^ respectively) or on both ends (RNA^dual-OH^). All of the extended ssRNA constructs have a higher predicted propensity to form local secondary structure (ΔG < -2.5 kcal.mol^-1^) relative to RNA^cognate^ (ΔG = 0.0 kcal.mol^-1^) (Figure 3D, Figure S1B). Incubation of dPPR10 with either RNA^5’-OH^ or RNA^3’-OH^ resulted in the formation of the high-FRET state indicative of ssRNA binding (Figure 3E) and consistent with ensemble-based SPR data (Figure S7A). Interestingly, only a small fraction of dPPR10 molecules occupied the high-FRET state when incubated in the presence of RNA^dual-OH^, indicating that the additional sequences flanking the cognate prevent (or slow down) its recognition and binding by dPPR10. The inability of dPPR10 to bind RNA^dual-OH^ could be either due to non-productive association with the additional sequence or exclusion from the cognate site, by, for example, RNA secondary structures. If either of these hypotheses are correct, we would expect preventing interaction between the extensions and the target site of dPPR10 to improve binding. Indeed, incubation of RNA^dual-OH^ duplexed with DNA primers annealed to the sequences flanking the cognate target site resulted in a stark increase in the fraction of molecules with high FRET efficiency to a level similar to that observed for RNA^cognate^ (Figure 3E, Figure S7B). Incubation of the DNA primer alone did not result in an increase in FRET efficiency; indeed, native-PAGE analysis confirmed that the addition of DNA primer serves only to inccrease the binding preference of PPR to duplexed RNA^dual-OH^ species (Figure S7C). To distinguish between association of dPPR10 with the extension sequences and association of the cognate target with the extension sequences as the explanation for the inhibition of binding, we prepared an ssRNA oligonucleotide of identical length to RNA^dual-OH^ (53 bp) but without predicted secondary structure (denoted as RNA^dual-OH^ ^(Δ2°)^). dPPR10 binds to this RNA as well as it binds to RNA^dual-OH^ duplexed with DNA primers (Figure 3E). This suggests that the key limitation to binding is not the length of the RNA species *per se*, nor association of the PPR with non-cognate sequence, but rather RNA secondary structure occluding the target site.

### Increased RNA transcript length and secondary structure propensity slows association to PPR

Using microfluidics, we determined the time taken until an increase in FRET was observed following injection of the RNA constructs into the flow cell containing dPPR10 (Figure 3F-G, additional traces in Figure S7D). As expected, RNA^cognate^ was the fastest to induce an increase in FRET (mean binding time of 10.6 ± 0.8 s), with RNA^5’-OH^ and RNA^3’-OH^ taking four times longer (41.3 ± 4.9 s and 43.4 ± 4.6 s, respectively) and RNA^dual-OH^ taking 5 times longer (53.0 ± 10.2 s) to cause an increase in FRET. However, when DNA primers were annealed to RNA^dual-OH^, dPPR10 was able to bind to the duplexed construct substantially faster with a mean binding time of 24.0 ± 2.2 s (i.e. two-fold that of RNA^cognate^, Figure 3G). dPPR10 was observed to bind to RNA^dual-OH^ ^(Δ2°)^ with a similar mean binding time (30.0 ± 3.0 s) compared to RNA^dual-OH^ duplexed to DNA primer, which suggests that the presence of flanking ssRNA (as opposed to PPR-inaccessible RNA:DNA duplexes) does not assist (but may delay) the dPPR10 protein in recognizing and binding to its cognate sequence within a longer RNA molecule. The slow binding kinetics of dPPR10 when incubated in the presence of all RNA extension constructs is supported by T_low-high_ residence times, which are similar to the corresponding binding times (Figure S7E). These data extends on the previous observation that the length of an RNA transcript affects association rates (e.g., truncated ssRNA species associate to dPPR10 faster than RNA^cognate^), even in the absence of RNA secondary structure.

Interestingly, for all ssRNA constructs the T_low-high_ FRET transitions occurred abruptly from one frame to the next (Figure 3H), which suggests that the longer T_low-high_ residence times are not due to some rate-limiting conformational rearrangement of dPPR10 (which would result in a gradual increase in FRET efficiency). Intriguingly, smFRET experiments demonstrate that the presence of 100-fold excess of non-target RNA (i.e., RNA^anti-cognate^) did not affect the rate or amount of dPPR10 association to RNA^cognate^ (Figure S7F-G), whereas non-target sequences that flank the cognate sequence (e.g., RNA^dual-OH^ ^(Δ2°)^) affect binding kinetics. These data indicate that the delay in dPPR10 conformational compression in the presence of longer sequences is not due to competition between the target and non-target sequences for dPPR10 recognition but is rather a topological issue, whereby dPPR10 likely has to associate with the target sequence in the correct orientation (i.e., 5’ to 3’ direction) without perturbation from non-target flanking sequences.

### dPPR10 does not ‘scan’ non-cognate sequences and binds directly to the cognate site

It is interesting that we observed a substantial binding delay of dPPR10 to RNA sequences that contain secondary structure but do not detect any conformational rearrangement of the PPRs prior to compaction. To determine if we were missing any intermediate bound states where dPPR10 had not undergone conformational changes, we set out to perform 3-colour smFRET experiments with fluorescently-labelled ssRNA to directly correlate ssRNA binding with dPPR10 conformational changes. To do so, immobilized dPPR10 was incubated with an AF488-labelled RNA^cognate^ (AF488-RNA^cognate^) and alternatively excited with a 532 nm laser (to monitor dPPR10 conformational changes using smFRET, denoted FRET^PPR^) and a 488 nm laser (to visualize the binding of labelled ssRNA) (Figure 4A). As expected, incubation of dPPR10 with AF488-RNA^cognate^ resulted in the formation of the high-FRET^PPR^ state (∼ 0.7) typical of ssRNA-bound dPPR10, indicating that the attachment of a fluorophore to RNA^cognate^ does not affect its ability to induce dPPR10 conformational changes (Figure 4B). Notably, the signal from the AF488-RNA molecule appeared in the same frame as the T_low-high_ FRET^PPR^ transition (Figure 4C), indicating that ssRNA binding immediately led to dPPR10 compaction. Further, excitation with the 488 nm laser led to a concurrent increase in the total fluorescence of all three dyes associated with the dPPR10-ssRNA system (i.e., AF488, Cy3 and AF647), indicating that AF488-RNA^cognate^ is participating in FRET with the fluorophores conjugated to dPPR10 (Figure 4C-D, Figure S8). Likewise, transitions to low-FRET^PPR^ states (i.e., < 0.5) were coincident with a loss of fluorescence from all dyes upon excitation at 488 nm (Figure 4C). Interestingly, similar results were observed for AF488-RNA^5’-OH^, where although AF488-RNA^5’-OH^ did not bind to dPPR10 as efficiently as the cognate (Figure 4B), the increases in FRET^PPR^ observed were directly correlated with an increase in dye fluorescence following excitation at 488 nm (Figure 4C-D). Notably, in the presence of either labelled ssRNA constructs, when dPPR10 exists within the low-FRET^PPR^ state (i.e., apo-dPPR10) the total fluorescence of all of the dyes was close to zero upon excitation at 488 nm; conversely, when dPPR10 resided in the high-FRET^PPR^ state (i.e., ssRNA-bound dPPR10) the total fluorescence of all the dyes was substantially higher (Figure 4E). Collectively, these data indicate that the association and dissociation of ssRNA to dPPR10 is directly correlated with its observed transitions to a conformationally compact or relaxed state, respectively. Furthermore, since a non-specific ssRNA-dPPR10 intermediate complex could not be observed, this data demonstrates that dPPR10 binds directly to the cognate sequence and has limited or no capacity to ‘scan’ ssRNA transcripts. Such a binding model provides a simple explanation as to why sequestration of cognate sequences within secondary structure elements can effectively prevent dPPR10 binding.

**Figure 4:**
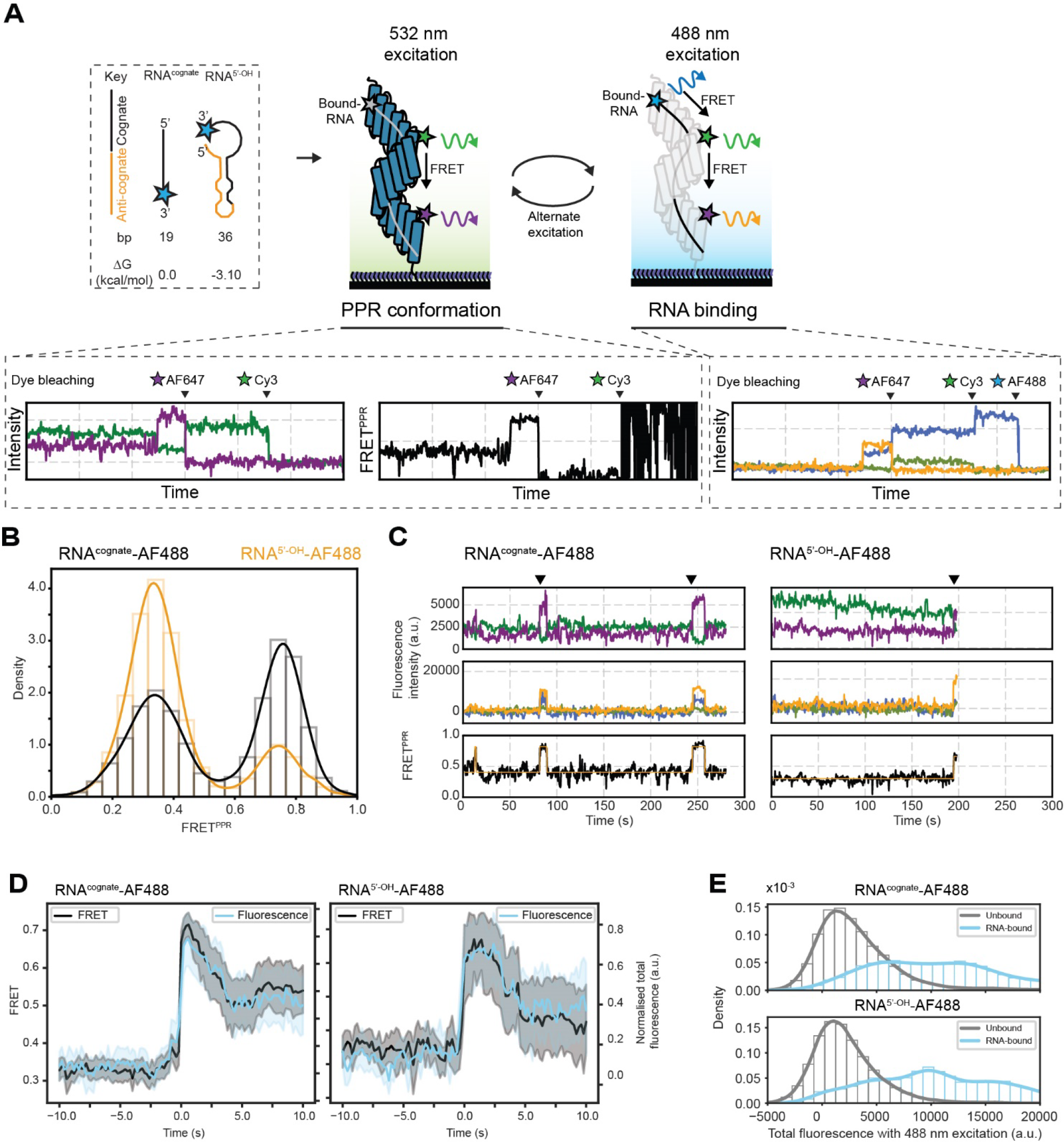
dPPR10 conformational compaction occurs only upon specific recognition and binding to the cognate sequence. (A) Schematic of the 3-colour smFRET experimental setup. Immobilized Cy3/AF647-labelled dPPR10 is incubated in the presence of AF488-RNA^cognate^ or AF488-RNA^5’-OH^ (20 nM) and alternatively excited with a 532 nm laser (to monitor changes in dPPR10 FRET) or 488 nm laser (to monitor association of labelled ssRNA to dPPR10) and the fluorescence emission of each dye is measured over time. For clarity, the point at which each of the dyes photobleaches during imaging is shown above each example trace. Example RNA molecules are not shown to scale. **(B)** FRET histogram of dPPR10 incubated in the presence of AF488-RNA^cognate^ or AF488-RNA^5’-OH^. Data is collated from at least 282 individual molecules. **(C)** Representative fluorescence intensity traces of dPPR10 and either AF488-FRET^cognate^ (*left*) or AF488-RNA^5’-OH^ (*right*) when excited with the 532 nm laser (*top*), the 488 nm laser (*middle*) and FRET^PPR^ trajectories upon excitation with the 532 nm laser (*bottom*). The arrows above the traces indicate events where an increase in FRET^PPR^ is correlated with the association of AF488-RNA. **(D)** Synchronized transition plots showing the PPR^FRET^ and normalised AF488-RNA^cognate^ (*left*) or AF488-RNA^5’-OH^ (*right*) total fluorescence immediately prior to and after a T_low-high_ transition (indicative of ssRNA binding). **(E)** The total fluorescence intensity of AF488, Cy3 and AF647 fluorophores upon 488 nm excitation depending on whether FRET^PPR^ is < 0.5 (*unbound*) or > 0.5 (*RNA-bound*) in the presence of either AF488-RNA^cognate^ (*top*) or AF488-RNA^5’-OH^ (*bottom*).

## Discussion

We present a crystal structure of a P-class designer PPR protein bound to a target RNA molecule and used this system to develop and characterize a robust PPR-based FRET sensor for RNA binding. We then interrogated the effects of RNA modifications (e.g., mismatches and secondary structure) on dPPR10 binding using single-molecule fluorescence approaches. We demonstrate that dPPR10 becomes conformationally compressed when bound to RNA, consistent with previous reports of other PPR proteins ^12,14^, and that this compaction is supported by contact of the RNA phosphate with Lys13 within each motif (Figure 1D, Figure S3E). Furthermore, smFRET and SPR data revealed that mismatches within the center of the binding site, but not the 3’ end, of the cognate sequence severely impairs association to dPPR10 and that a minimum of 10 PPR-nucleobase contacts are required to induce binding to dPPR10. Notably, two- and three-colour smFRET highlights that RNA secondary structure reduces the accessibility of binding sites to dPPR10.

It has been proposed that PPR10 regulates the translation of *atpH* by preventing the formation of secondary structure directly upstream of the ribosome binding site ^7^, and prevents nuclease attack ^5^, with upstream binding of start codons also observed for other PPR proteins ^42,43^. Paradoxically, however, the binding of PPR to RNA sequences containing secondary structure elements is impaired *in vitro* ^44^, which begs the question as to how PPR deals with such challenges. The data reported in this work supports a model whereby the presence of RNA secondary structure slows or prevents dPPR10 binding by sequestering the cognate sequence such that it cannot be readily accessed by dPPR10, which itself has no ability to bind to these structures. Our data, as well as others, indicate binding to secondary structure elements does not occur since PPR motifs make direct contact with the face of the RNA nucleobase ^12,45^. Indeed, even when portions of the cognate sequences are accessible within modelled RNA structures ^46^, binding to these sequences remained impaired compared to RNA^cognate^; this data suggests that secondary structures introduce entropic constraints that may interfere with key PPR-ssRNA contacts (i.e., Lys13, amino acids at position 5 and 35) that enable initial recognition and binding.

As such, the observed delay in dPPR10 binding to ssRNA sequences containing predicted secondary structure is likely due to the kinetics of ssRNA secondary structure formation and disappearance. The finding that even relatively weak ssRNA secondary structures (ΔG = ∼ -2.5 kcal.mol^-1^) have a substantial effect on the ability of dPPR10 to bind to target sequences is somewhat surprising considering that certain PPR proteins, including PPR10, are known to bind at sites *in vivo* that can form hairpin structures ^47,48^. Notably, PPR10 and PPR53 binds to sites *in vivo* that have similar or higher secondary structure stability (ΔG = -2.8 or -8.0 kcal.mol^-1^, respectively) compared to the ssRNA variants used in this work ^7,47^. Indeed, the stability of these secondary structures are likely to be substantially higher *in vivo* compared to *in vitro* due to macromolecular crowding, which is known to promote folded and compact ssRNA structures ^49,50^. The reduced ability of dPPR10 to bind to sites containing secondary structure appears to be in conflict with its proposed mechanism of translation upregulation; it is likely that PPR proteins performs this role in cooperation with other proteins *in vivo*, such as RNA helicases, chaperones or sRNAs ^51^, that may temporarily resolve secondary structure elements to facilitate stable PPR binding.

Despite the well-established role of PPR proteins in RNA regulation, the question as to how they find their target sequences in the complex cellular milleu within biologically relevant timescales remains an under- explored area of study. Two models are possible; either a PPR protein binds to sequences upstream or downstream of the target sequence and ‘scans’ the ssRNA transcript until the target is encountered (2- dimensional search, fast association [known as facilitated diffusion]) or it opportunistically binds to exposed target sequences (3-dimensional search, slow association). The data presented in this study supports the latter model, since we do not observe any transient association of dPPR10 to longer ssRNA sequences prior to conformational compaction as observed using 3-colour FRET. Furthermore, RNA^anti-cognate^ did not show any capacity to bind dPPR10, even transiently as determined by SPR and smFRET, which again suggests that dPPR10 cannot associate with non-target sequences. For proteins restricted to 3D-diffusion-dependent searching, high protein concentrations are favoured since the probability of direct interaction with the target sequence is increased ^52^; conversely, excess non-target sequences reduce the probability of direct protein-target collisions and decreases association rates. Interestingly, direct binding to sequence-specific sequences (facilitated primarily by 3D-diffusion) by nucleic-acid-binding proteins has been experimentally observed even in the presence of massive excess of non-target flanking sequences ^53^. It was proposed that direct binding is enhanced by unusual nucleic-acid structural features close to the target sequence, which improves protein accessibility or specific-binding propensity ^53^. It is tempting to speculate that local RNA structures also act to improve the probability of specific PPR binding; for example, RNA secondary structures disproportionally decrease the number of single-stranded, non-specific target sequences (owing to their increased abundance), which biases accessible binding sites towards target sequences and reduces the effective ‘search space’ for productive PPR binding.

We propose the following mechanism of dPPR10 recognition and binding to ssRNA (Figure 5). First, secondary structures sequester the cognate nucleobases from PPR motifs, preventing binding (Figure 3D-H, Figure 4B-C). Transient exposure of the cognate sequence, the rate and propensity of which is primarily dictated by the inherent kinetics of RNA secondary structure and formation *in vitro* and possibly by other cofactors *in vivo*, allows cognate recogntion by PPR. Second, the initial contact of dPPR10 to RNA is via direct association of the first 2-3 PPR motifs with the cognate nucleobases and does not require ‘scanning’ of non-cognate sequences to do so. This is evidenced by several key findings, including (i) initial recognition of RNA by dPPR10 cannot be via non-specific lysine interactions with the RNA backbone, since the presence of excess non-target RNA species does not affect the rate of or binding to cognate (Figure S7F-G); (ii) dPPR10 does not associate at all to non-cognate sequences (Figure 2D, Figure 4B-C, Figure S7F-G); and (iii) dPPR10 binds to RNA^dual-OH^ duplexed to DNA primer at similar rates as RNA^dual-OH^ ^(Δ2°)^, which indicates that there is no improved capacity of dPPR10 to search for the cognate if it is flanked by ssRNA vs duplexed nucleic acid (Figure 3D-H). Third, establishing a minimum of two PPR-nucleobase coordinates the interaction between Lys13 of the first PPR motif (*p*) with the phosphate backbone of a downstream nucleobase (*b+2*), which results in charge neutralisation of the lysine that annuls its repulsive force with Lys13 in the adjacent PPR module. Consequently, the adjacent module can partially contract to cooperatively establish contacts with the next nucleotide within the ssRNA sequence; this cycle of charge neutralization and motif compaction then propagates throughout the entire protein. Mutagenesis of Lys13 completely abolishes binding of PPR to ssRNA ^12^, which supports our proposed model that a combination of specific amino-acid/nucleobase and Lys13-phosphate interactions are essential in overcoming the energy barrier for conformational rearrangements and stable ssRNA binding to dPPR10. Disruption of PPR motif and nucleobase interactions (e.g., by double mismatches) at early positions can disrupt the propogation of motif compaction upon binding (Figure 2F-G) or promote erronous motif expansion during dissociation (Figure 2G-H), which substantially reduces the *K*_D_. Notably, combined incubation of truncation variants does not restore the conformational compaction of dPPR10 to levels observed for RNA^cognate^, indicating that stable binding occurs only when a single continuous RNA molecule is present to coordinate the propagation of conformational changes throughout the entire protein.

**Figure 5:**
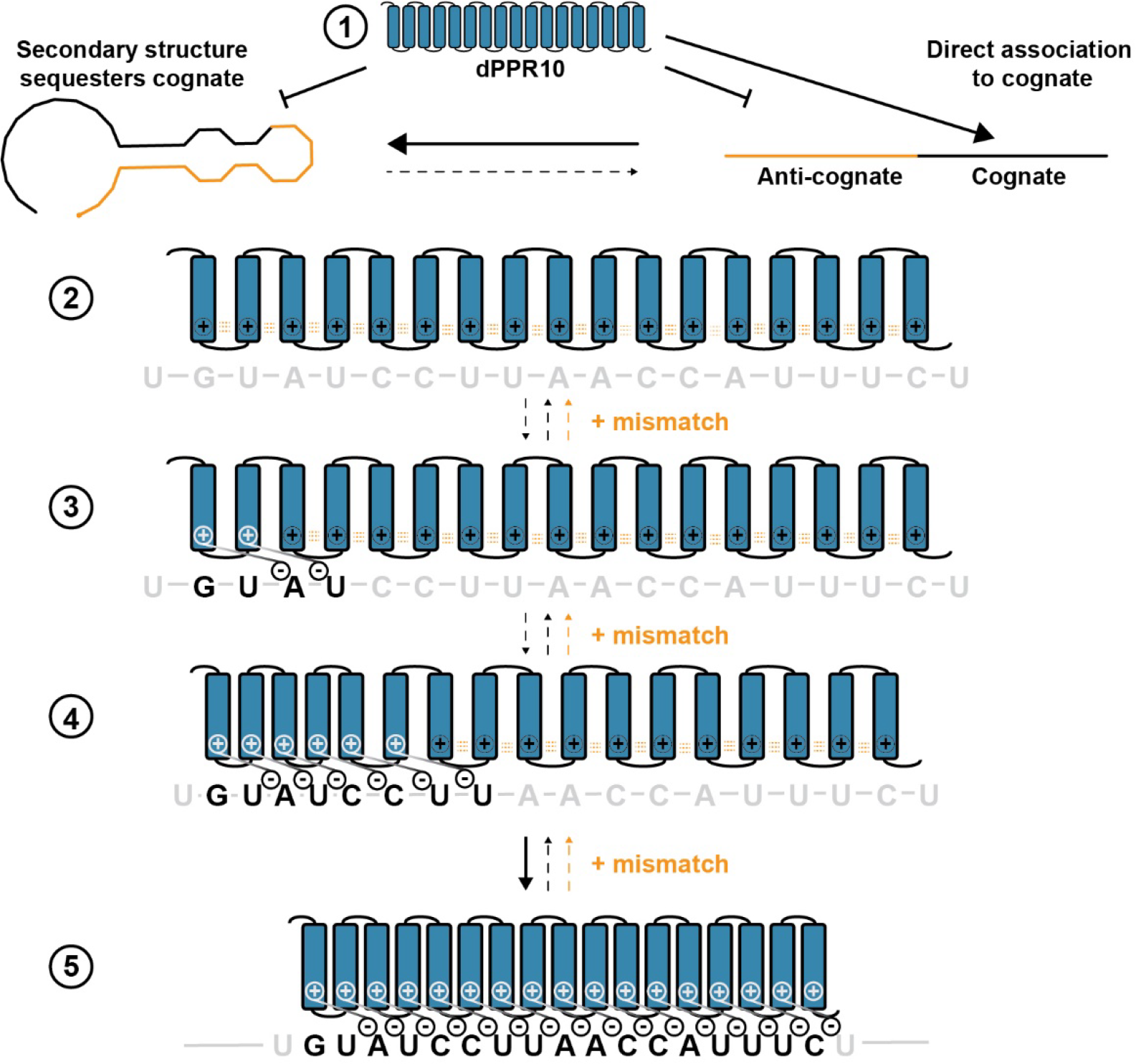
Binding mechanism of dPPR10 to ssRNA. The presence of stable ssRNA secondary structure sequesters the cognate sequence, which prevents the association of dPPR10. If the stability of ssRNA secondary structure is sufficiently low, it can stochastically sample folded and unfolded configurations to which the dPPR10 will opportunistically associate with the transiently exposed cognate sequence (but not non-target ssRNA sequence). **(2)** apo-dPPR10 exists in a conformationally expanded state, driven by coulombic repulsion (*dotted orange lines*) between Lys13 residues in adjacent PPR motifs. **(3)** Initial PPR-nucleobase recognition at the 5’/N-terminal end is made (*black*), which stabilizes an electrostatic bridge between Lys13 in PPR module p and the phosphate moiety in nucleotide *b+2*. **(4)** Charge neutralisation caused bt the Lys13-phosphate interaction reduces the coloumbolic repulsion that is common in the apo- dPPR10 conformation, resulting in local compression of PPR motifs that enables additional PPR-nucleobase and Lys13- phosphate interactions to be established. **(5)** At this stage, the charge neutralisation effect becomes strongly favoured (due to the established multivalent interactions), which then propogates throughout the dPPR10 structure for stable binding. The presence of mismatches (or truncations) can significantly increase the reverse reaction (*shown in orange*).

RNA- or DNA-binding proteins typically bind to their target sequences faster than random, three-dimensional diffusion would predict ^54,55^; this has been proposed to occur via facilitated difussion, whereby slow, non- specific association of protein to nucleic acid (3D diffusion dependent) is followed by faster two-dimensional scanning (via hopping or sliding) until the cognate sequence is encountered ^52^. Since dPPR10 does not associate appreciably with non-cognate RNA, two-dimensional scanning is restricted and is instead more dependent on 3D diffusion. The kinetic constraints imposed by random 3D diffusion may explain the longstanding maxim and associated conundrum of why PPR proteins are abundant in organelles, but almost unknown outside them (i.e. in the cytosol and nucleus). Other functionally analogous but non-homologous RNA binding protein families (e.g., mTERF proteins ^56^, HPT proteins ^57^, OPR proteins ^58^) show a similar restriction to organelle compartments, perhaps for the same reason. Collectively, these findings have significant implications for the design of synthetic PPR proteins. Careful consideration must be given to the local binding environment (e.g., primary and secondary structure of RNA target) within the global transcriptome. As reported by others ^18^, dPPR10 binding appears to be relatively tolerant to mismatches toward the 3’ end compared to those within the center of the PPR binding site or 5’ end; however, single mismatches dramatically increase dissociation rates (irrespective of sequence position). While seemingly a negative feature, transient interactions of a PPR with an RNA transcript may be beneficial in some applications (i.e., to track localization of transcripts *in vivo* without perturbing function or transcript lifetime) but could be detrimental to others if stable binding is preferred (e.g., if attempting to modulate gene expression). Furthermore, PPR binding is surprisingly affected (or even completely abolished) by even weak secondary structure elements (ΔG = ∼ -2.5 kcal.mol^-1^); as such, the design of synthetic PPRs should be targeted to RNA sequences within an RNA transcript that contains low predicted secondary structure. Finally, our data indicates that dPPR10 has a limited capacity to ‘search’ ssRNA transcripts for its target sequence. As such, this may have cautionary implications for the design of and use of synthetic PPRs in nuclear and cytoplasmic settings, restricting the use of dPPRs within organelle compartments, unless other approaches can be developed to increase the scanning speed of PPRs for their targets (e.g., fusing an RNA helicase to a PPR module).

## Acknowledgments

This research was undertaken in part using the MX2 beamline at the Australian Synchrotron, part of ANSTO, and made use of the Australian Cancer Research Foundation (ACRF) detector. We would also like to thank the staff at Molecular Horizons (UOW) for their technical and administrative support.

## Author contributions

Research concept was developed by BJ, BP, TB, MA, IS, AvO and CSB. Experiments were designed by all authors. Experiments and data analysis was carried out by NM, BJ, BP, JS, CSB and SJ. The manuscript was written by NM, BJ, MA, IS, AvO and CSB.

## Supplementary Tables

**Table S1:**
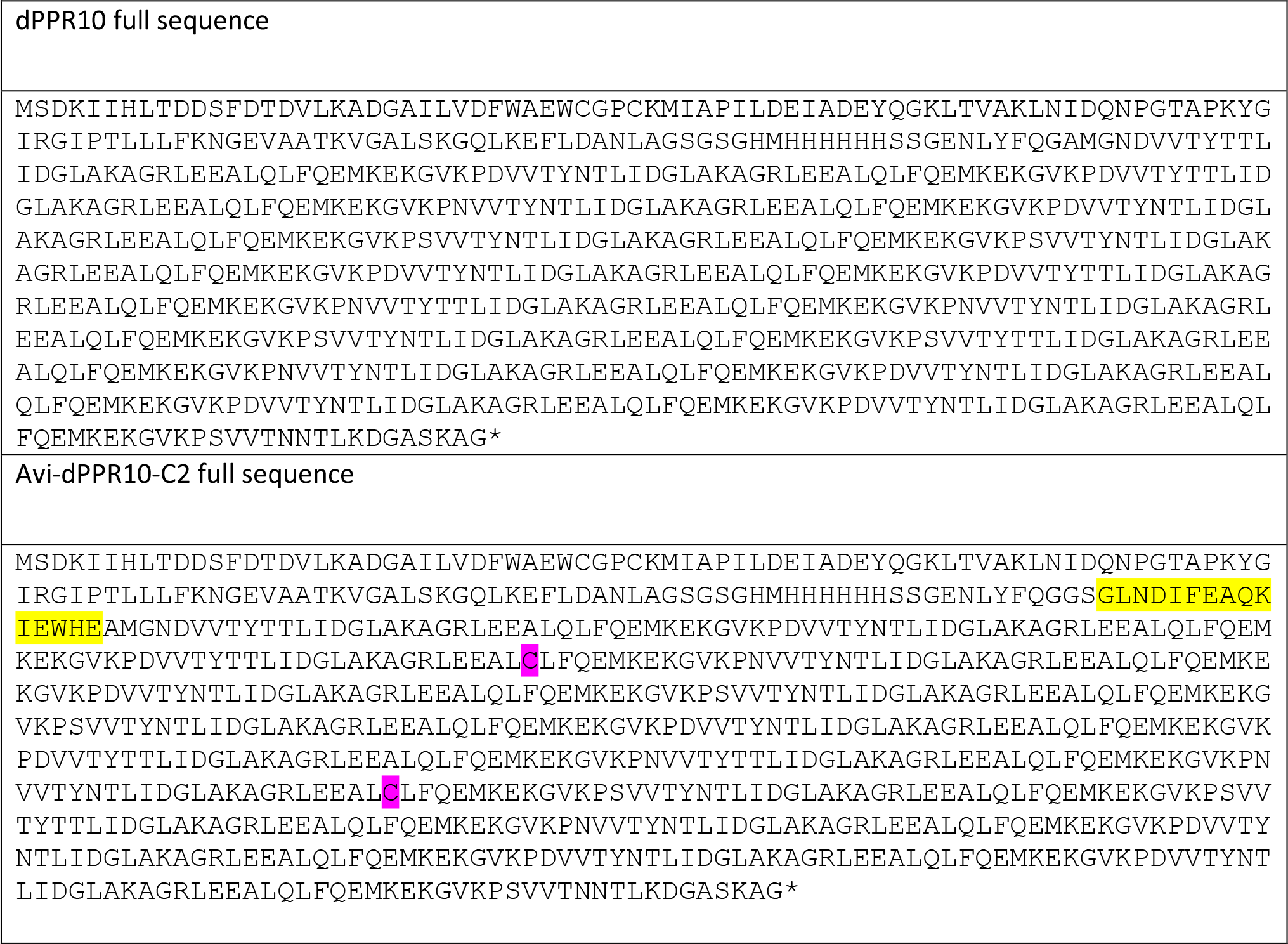
Complete protein sequences used in this work. Yellow and pink denotes the position of the AviTag sequence and the position of cysteines for labelling, respectively.

**Table S2:** Crystallographic data and refinement statistics. Numbers in parenthesis correspond to the highest resolution shell.

|  |  |
| --- | --- |
| **Data collection** |  |
| **Space group** | *P1* |
| **Unit cell dimensions** |  |
| **a, b, c (Å)** | 43.34, 51.75, 51.97 |
| **$\alpha, \beta, \gamma$ (°)** | 118.1, 97.2, 96.0 |
| **Wavelength (Å)** | 0.9537 |
| **Resolution (Å)** | 44.92-2.01 (2.06-2.01) |
| **R<sub>merge</sub> (%)** | 5.1 (22.8) |
| ***I*/σ*I*** | 9.0 (2.1) |
| **CC <sub>1/2</sub>** | 0.995 (0.786) |
| **Completeness (%)** | 97 (90.8) |
| **Redundancy** | 2.0 (1.6) |
| **Wilson B (Å<sup>2</sup>)** | 26.9 |
| **Refinement** |  |
| **Resolution (Å)** | 44.92 – 2.00 |
| **No. reflections** | 23919 |
| **R<sub>work</sub>/R<sub>free</sub>** | 18.2/24.7 |
| **No. Atoms** | 10755 |
| **Protein** | 9773 |
| **RNA** | 768 |
| **Water** | 214 |
| **Average B-factor (Å<sup>2</sup>)** | 32.2 |
| **R.m.s deviations:** |  |
| **Bond lengths ( Å )** | 0.014 |
| **Bond angles (°)** | 1.6 |
| **Ramachandran analysis** |  |
| **Favored (%)** | 96.0 |
| **Allowed (%)** | 4.0 |
| **Outliers (%)** | 0.0 |

## Supplementary figures

**Figure S1:**
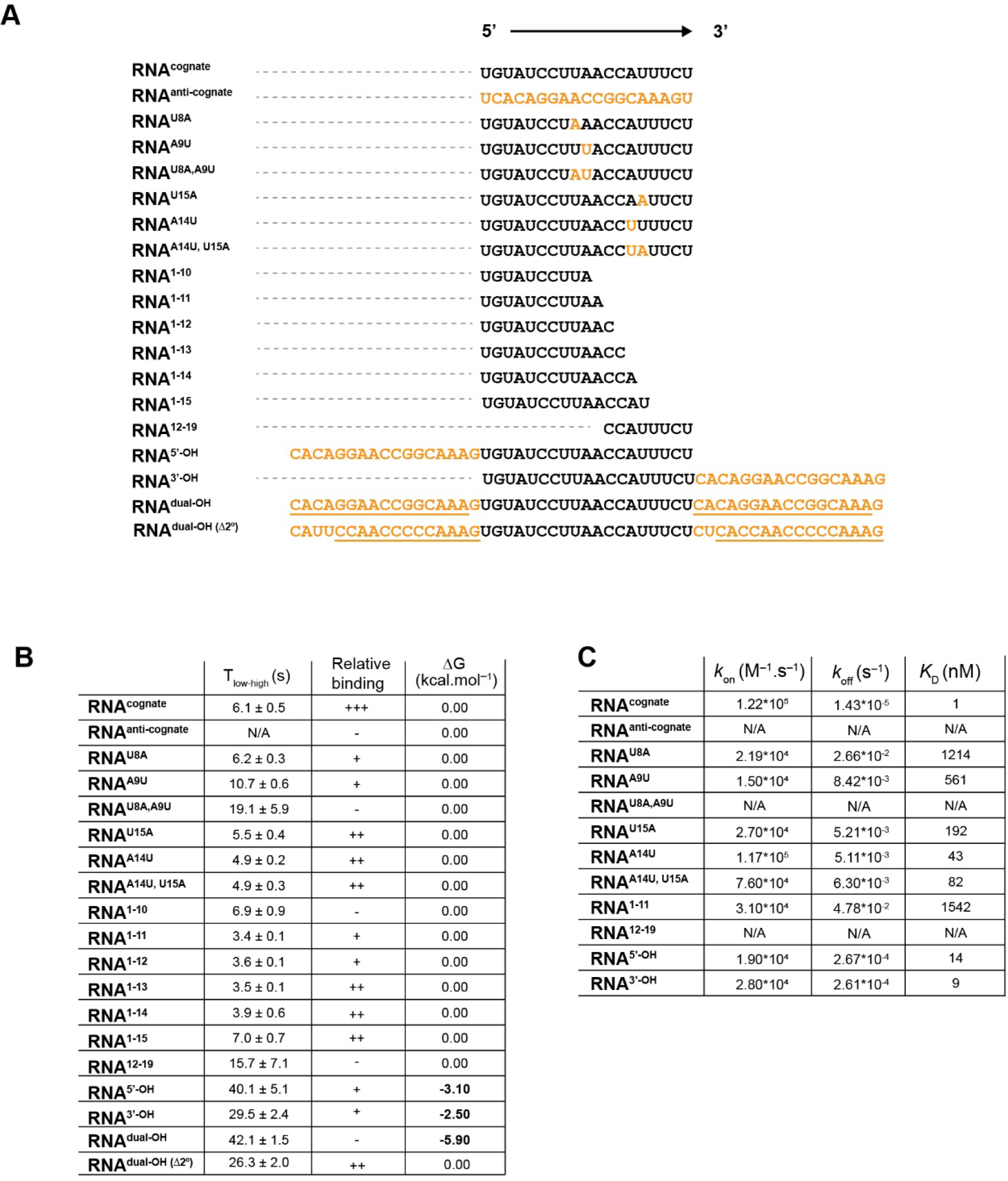
Schematic and sequence of each ssRNA oligonucleotide used in this work with summarised kinetic information. (A) The identity and sequence of each ssRNA oligonucleotide is shown, with modifications to RNA^cognate^ (*orange*) and position of complementarity with DNA primers (*underlined*) denoted. **(B)** The T_low-high_ residence times (indicative of ssRNA binding) determined for each ssRNA variant as determined by smFRET, reported as the mean ± SEM. The relative binding efficiency (fraction of molecules in the high FRET state in the FRET efficiency histograms) and propensity of each ssRNA to form secondary structure is also shown. **(C)** Association and dissociation rates of various ssRNAs, with calculated *K_D_*, as determined using SPR.

**Figure S2:**
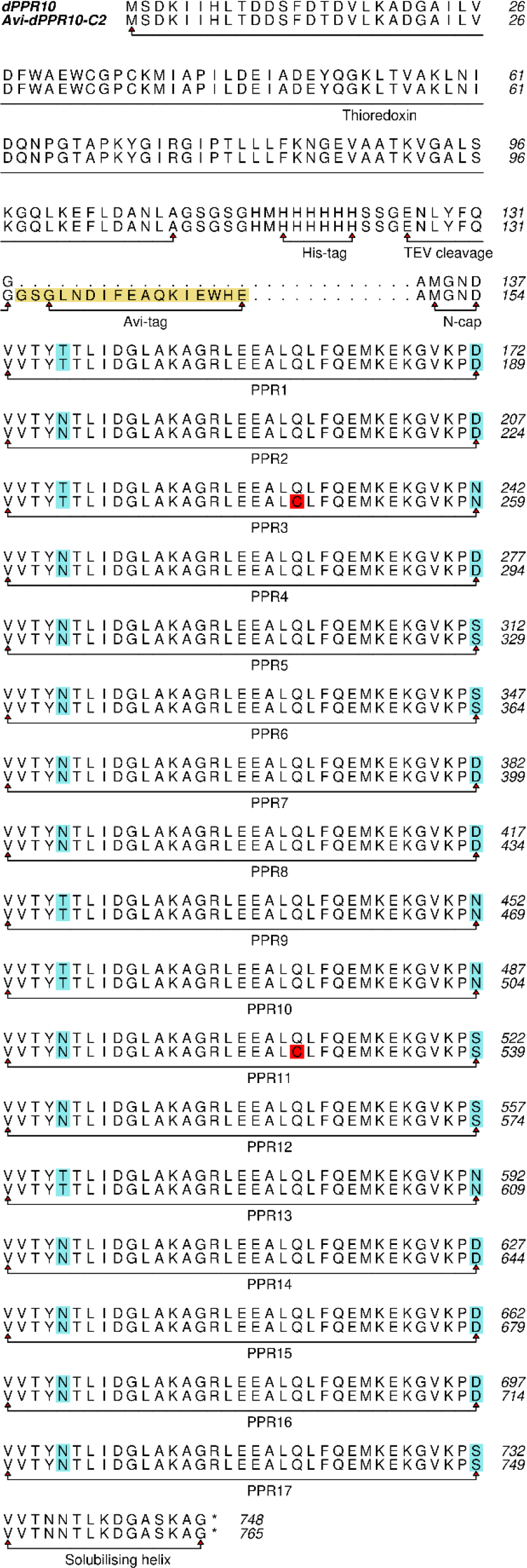
Details of protein sequences. Notable features are labelled or coloured (red indicates positions of cysteines introduced for labelling; cyan indicates RNA-specificity changes; yellow indicates the Avi-tag).

**Figure S3:**
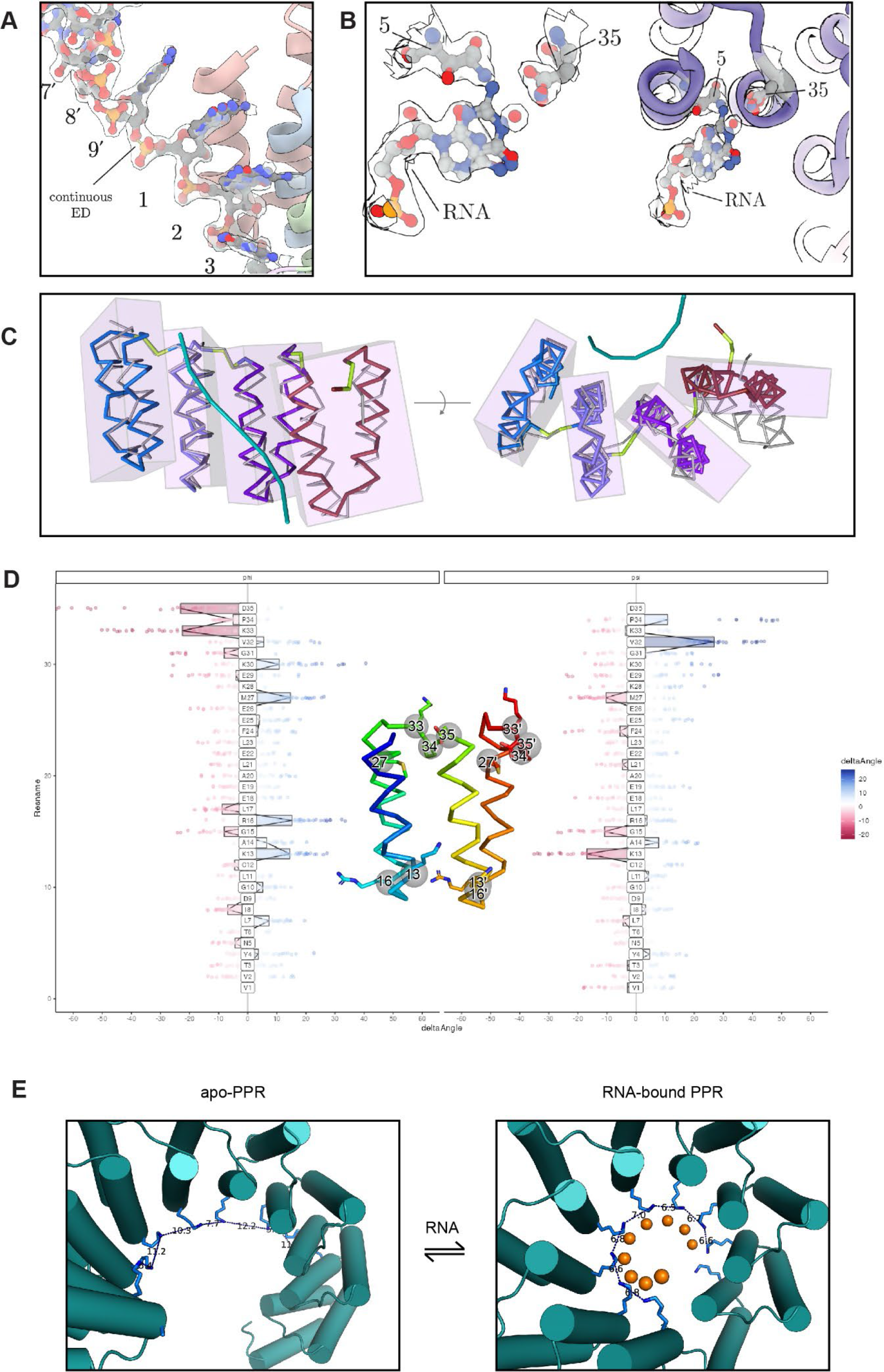
**Structural features of PPR-protein:RNA binding**. **(A)** Helical disorder in the structure of dPPR10:RNA complex results in fully connected eletron density between unit cells, and **(B)** microheterogeneity at the specific positions where nucleobases interact with residues 5 and 35. The structure otherwise has excellent geometry and electron density, representing an indeterminately long protein binding to an equally long RNA. **(C)** PPR repeats act as relatively rigid units (enclosed in transparent boxes) with a hinge region (green ribbon) between repeats, as analysed with DYNDOM. **(D)** Backbone geometry analysis (difference φ and ψ values at each position of a PPR repeat in the presence or absence of RNA, averaged across all repeats) reveals differences in protein conformation at only a small number of positions between apo- and ssRNA-bound structures (highlighted in inset), mainly corresponding to the hinge region. **(E)** Structural rearrangment in the presence of polyanionic RNA allow conserved cationic lysine sidechains to move closer together, a feature which may relate protein:RNA affinity to conformational change on binding.

**Figure S4:**
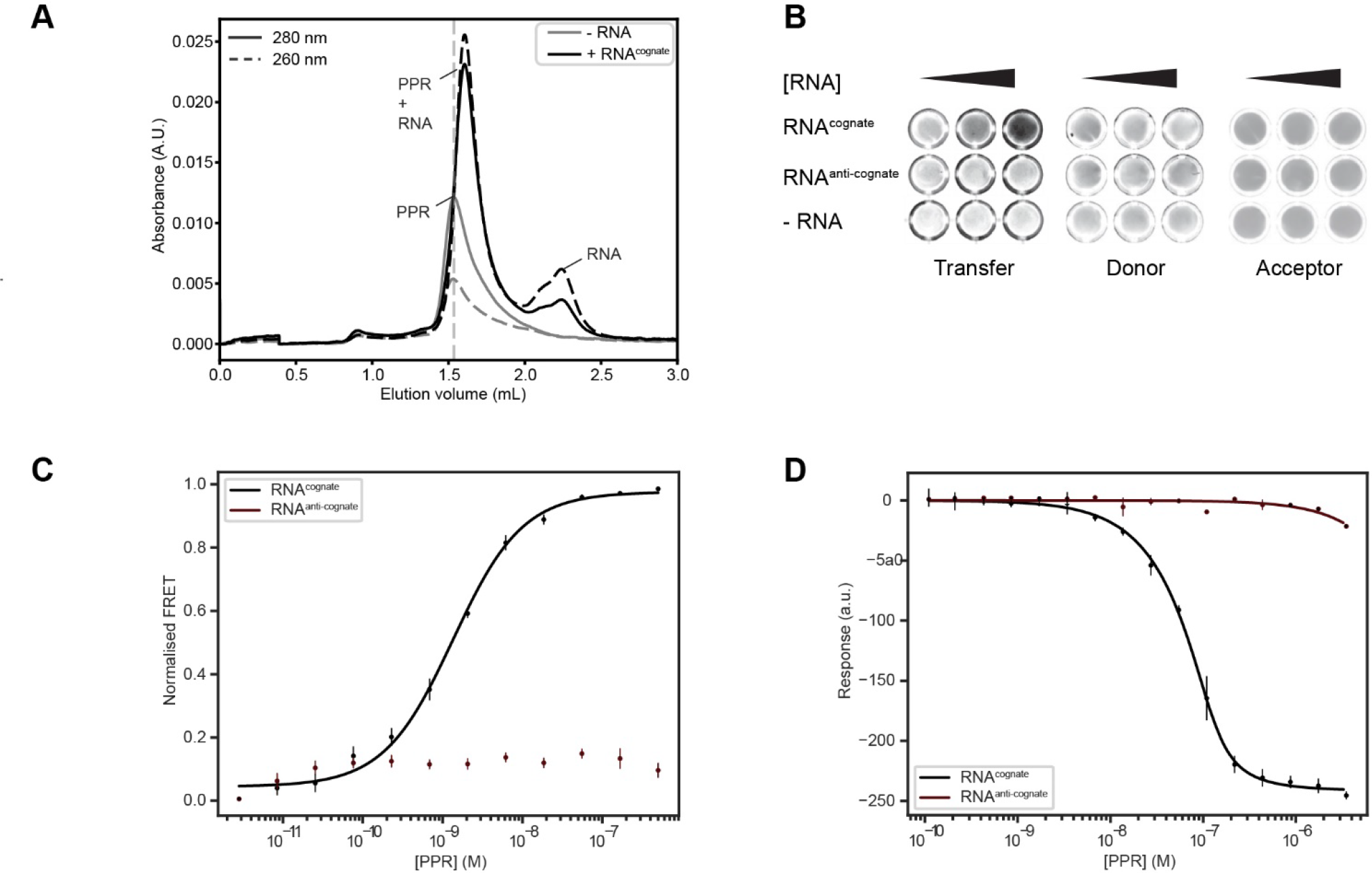
The conformational compaction of dPPR10 upon binding to ssRNA can be monitored in solution. **(A)** dPPR10 (15 µM) in the absence or presence of RNA^cognate^ (18 µM) was analysed using analytical size exclusion chromatography (Superdex 200) revealing shift to higher elution volumes and the relative decrease in hydrodynamic radius upon the addition of ssRNA. **(B)** Representative wells of a fluorescently imaged 96-well microtitre plate, showing the increase in fluorescence of the acceptor dye when exciting the donor in the presence of increasing concentrations of RNA^cognate^ (0.01 – 1000 nM). **(C)** Calculated FRET efficiency of labelled dPPR10 protein in the presence of either RNA^cognate^ or RNA^anti-cognate^, calculated from the bulk-FRET plate assay in (C). The *K*_D_ for RNA^cognate^ was calculated to be 1.2 ± 0.2 nM (average of 3 technical replicates). FRET was normalised to the maximum measured value for dPPR10 in the presence of RNA^cognate^. **(D)** Microscale thermophoresis of the fluorescently labelled dPPR10 protein, with RNA^cognate^ and RNA^anti-cognate^, showing a calculated binding affinity of 9.7 ± 1.8 nM (average of 3 technical replicates).

**Figure S5:**
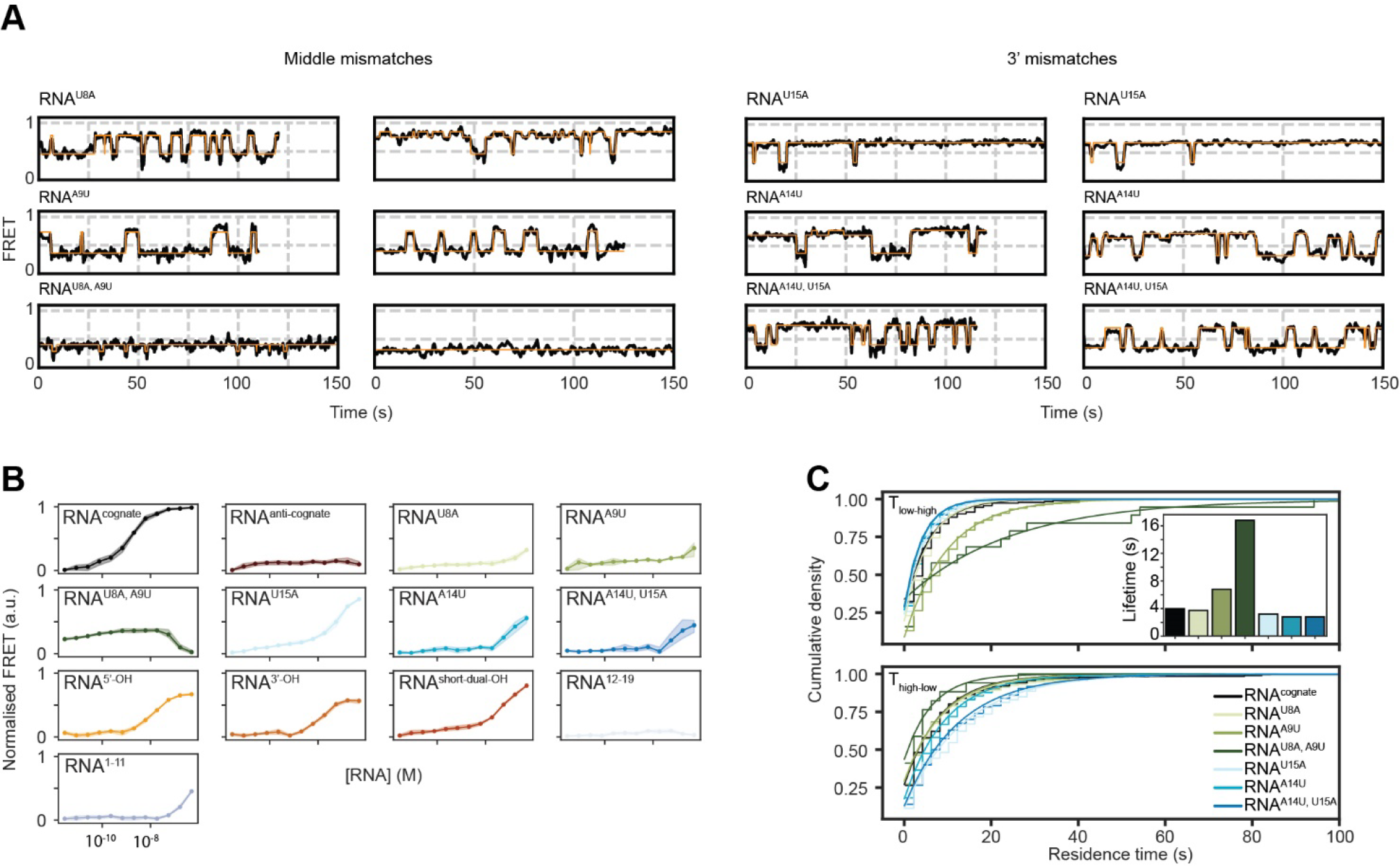
dPPR10 tolerates mismatches toward the 3’ end of ssRNA but not when mismatches are located centrally in the PPR binding site. (A) Representative FRET trajectories from individual dPPR10 molecules in the presence of the indicated ssRNA oligo containing mismatches within the dPPR10 region probed by FRET (1 µM, *left*) or at the 3’ end (1 µM, *right*). Orange lines represent the fit from the HMM. **(B)** Ensemble-based FRET titration plots of dPPR10 in the presence of various ssRNA constructs used throughout this study. **(C)** Cumulative histogram of the residence time of different transition classes. Data is fit with a one-phase association curve. For T_low-high_ residence times, the lifetimes calculated from the fit are shown.

**Figure S6:**
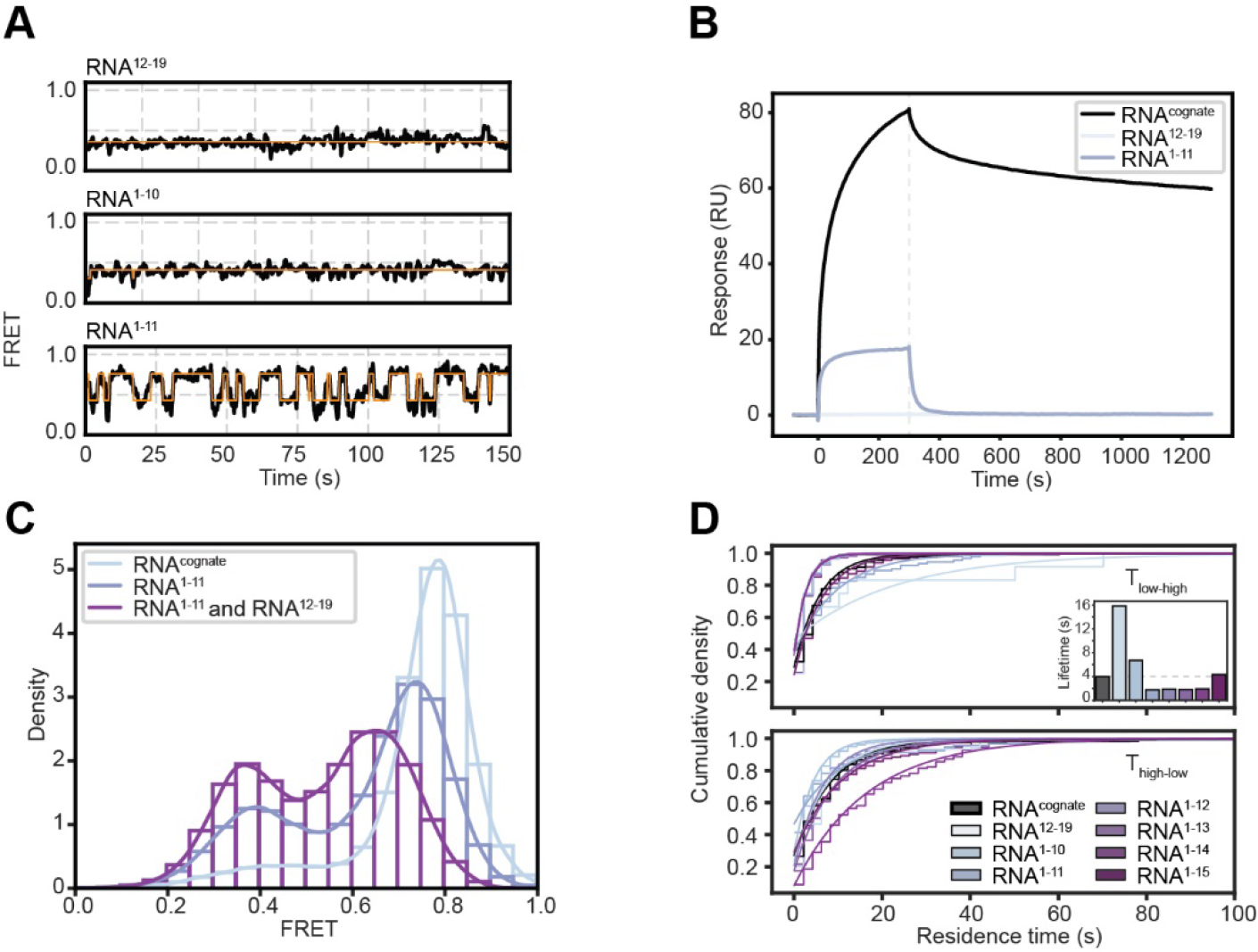
A minimum of 10 PPR-nucleobase contacts are required to induce conformational compaction of dPPR10. (A) Representative FRET trajectories from individual dPPR10 molecules in the presence of the indicated ssRNA oligo truncations (1 µM). Orange lines represent the fit from the Hidden Markov Model (HMM). **(B)** SPR association and dissociation curves of RNA^cognate^, RNA^1–11^ and RNA^12–19^ binding to dPPR10. **(C)** FRET histogram of dPPR10 incubated in the presence of RNA^cognate^ (1 µM), RNA^1–11^ (1 µM), or a combination of RNA^1–11^ and RNA^12–19^ (1 µM each). **(D)** Cumulative histogram of the residence time of different transition classes. Data is fit with a one-phase association curve. For T_low-high_ residence times, the lifetimes calculated from the fit are shown, with the dotted line indicating the value for RNA^cognate^ (*inset*).

**Figure S7:**
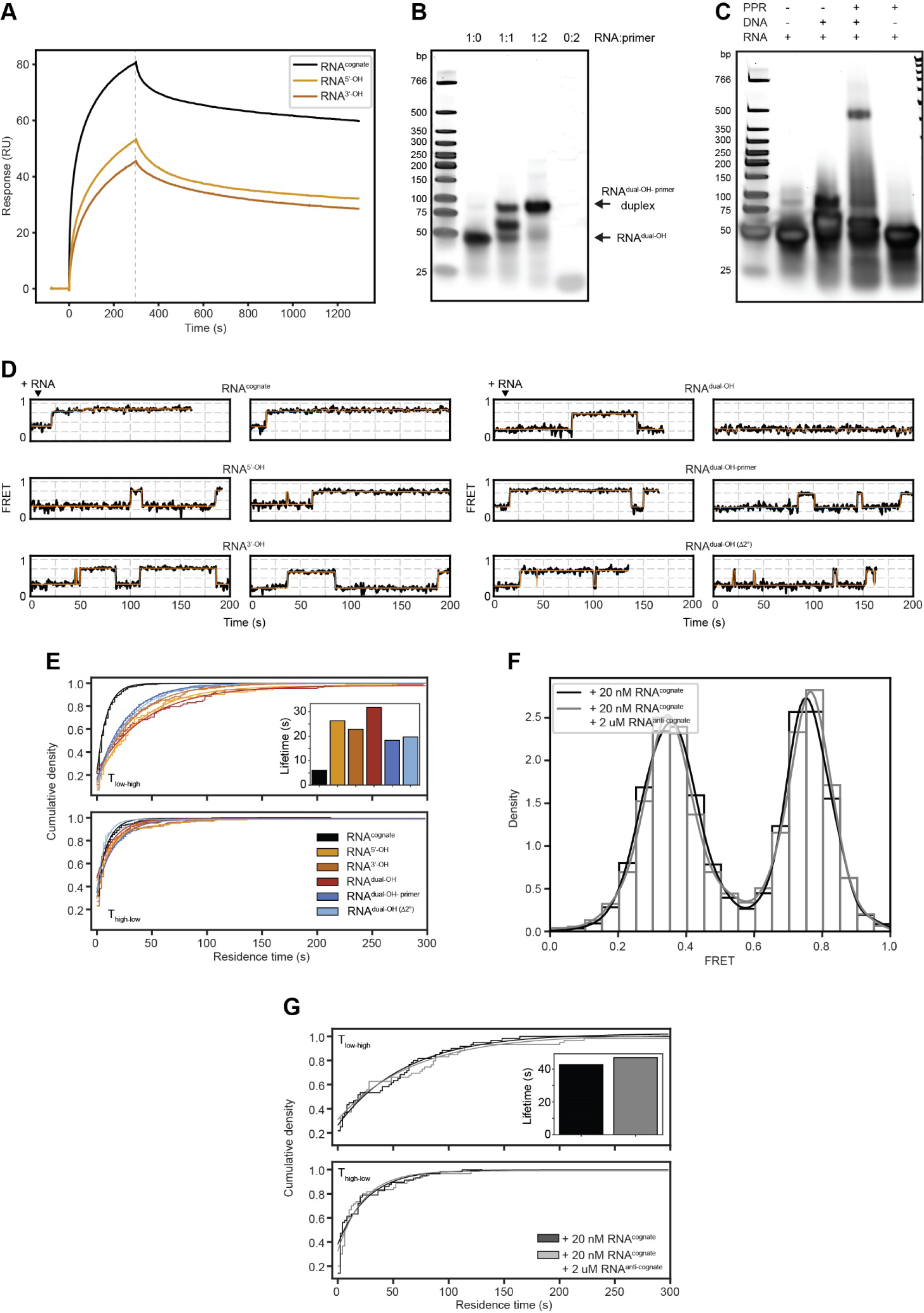
RNA secondary structure, oligonnucleotide length and the presence of flanking non-target sequences delays the binding of cognate ssRNA to dPPR10. (A) Surface plasmon resonance (SPR) traces of the indicated ssRNA constructs (1 µM) binding to immobilized dPPR10. Association of ssRNA to dPPR10 is initiated at t = 0 and proceeds for 300 s. Dissociation curves are generated following removal of ssRNA from solution. **(B)** Native SDS-PAGE gel of RNA^dual-OH^ (20 µM) following PCR melting and annealing in the absence or presence of DNA primer (20 or 40 µM, 1:1 or 1:2 molar ratio). Quick-load low molecular weight markers (in bp) are shown. **(C)** Native SDS-PAGE gel of RNA^dual-OH^ (1 µM) that had been incubated for 1 h at 4°C in the absence or presence of dPPR10 (1 µM) supplemented with or without DNA primer (2 µM). The combination of each component within each reaction is indicated above the lane. (**D**) Representative FRET trajectories of dPPR10 upon the injection of the indicated ssRNA (1 µM) after 10 s (indicated above the traces). **(E)** Cumulative histogram of the residence time of different transition classes. Data is fit with a one-phase association curve. For T_low-high_ residence times, the lifetimes calculated from the fit are shown. **(F)** FRET histogram of dPPR10 incubated in the presence of RNA^cognate^ (20 nM) supplemented with or without a 100-fold excess of RNA^anti-cognate^ (2 µM). Data is collated from at least 425 individual molecules. **(G)** Cumulative histogram of the residence time of different transition classes from treatments in which dPPR10 was incubated with RNA^cognate^ (20 nM) in the absence or presence of RNA^anti-cognate^ (2 µM). Data is fit with a one-phase association curve. For T_low-high_ residence times, the lifetimes calculated from the fit are shown.

**Figure S8:**
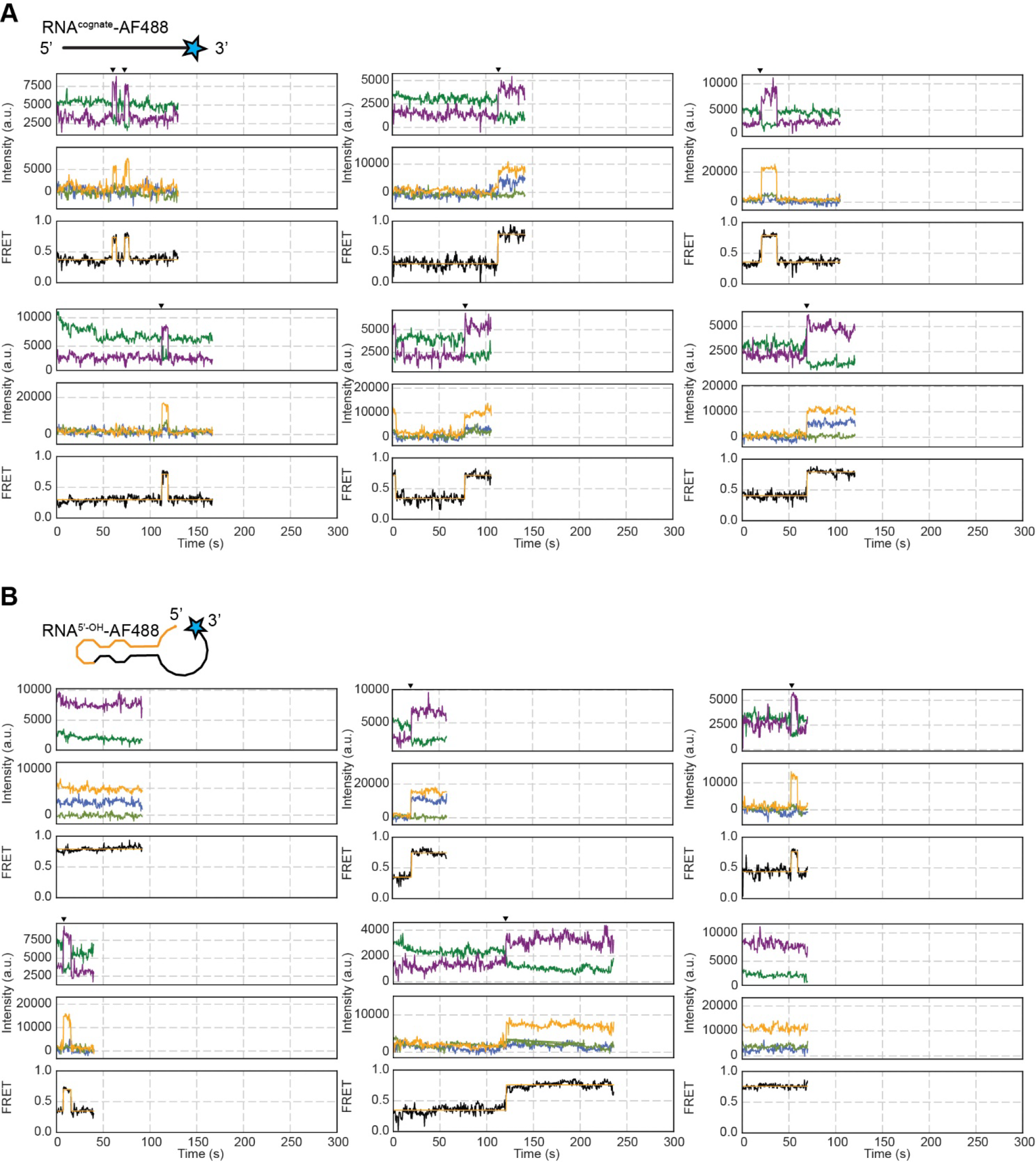
Representative fluorescence intensity and FRET^PPR^ trajectories from 3-colour smFRET experiments. dPPR10 was incubated in the presence of AF488-labelled **(A)** RNA^cognate^ (20 nM) or **(B)** RNA^5’-OH^ (20 nM) and the fluorescence from AF488, Cy3 and AF647 fluorophores was measured when excited with the 532 nm laser (*top*) or the 488 nm laser (*middle*). FRET^PPR^ trajectories upon excitation with the 532 nm laser (*bottom*) is also shown. Arrows indicate where increases in FRET^PPR^ are correlated with an increase in fluorescence following excitation at 488 nm.

